# Pathogenic human mitochondrial tRNA variants impair RNA processing by compromising 5′ leader removal

**DOI:** 10.64898/2026.03.25.714317

**Authors:** Jubilee H. Muñozvilla, Avery Ontiveros, Tatiana V. Mishanina

## Abstract

Human mitochondrial genome (mtDNA) encodes multiple proteins in the oxidative phosphorylation complexes as well as the ribosomal and transfer RNAs (tRNAs) needed for *in situ* translation. These genes are transcribed from only three promoters, producing polycistronic transcripts that are co-transcriptionally cleaved by mitochondrial RNase enzymes to release majority of individual gene products. tRNAs separate many of these genes and are thought to serve as “punctuation” marks that enable RNase recognition, binding, and hydrolysis of the 5′ “leader” and 3′ “trailer” sequences flanking the tRNA. Mutations in the tRNA genes dominate the mtDNA-linked mitochondrial pathologies; yet a systematic study of the impact of tRNA sequence variation on the RNase-catalyzed processing is lacking. Here, we employed human mitochondrial tRNA^Tyr^ as a model system to dissect the effect of tRNA variants on the *in vitro* 5′ leader and 3′ trailer hydrolysis. We found that nucleotide variations located near the catalytic interfaces – particularly within or near the tRNA acceptor stem – showed the strongest defects in 5′ processing and prevented release of the downstream tRNA in a tRNA cluster where multiple tRNAs are transcribed in tandem. This work provides mechanistic insight into how mutations disrupt coordinated mitochondrial tRNA processing and establish a framework for predicting variant effects based on their structural position relative to the processing enzymes.

## INTRODUCTION

Human mitochondria contain their own DNA (mtDNA), which encodes 13 essential protein subunits of the oxidative phosphorylation (OXPHOS) complexes that drive the synthesis of the majority of cellular adenosine triphosphate (ATP). The mtDNA-encoded genes are transcribed by a single mitochondrial RNA polymerase (POLRMT in humans) from three promoters, each producing a ∼16,600 nucleotide-long polycistronic transcript containing protein-coding sequences arranged in tandem (Fig. 1A). Processing of this primary transcript requires precise excision of intervening mitochondrial (mt) tRNAs by the endonucleases RNase P and RNase Z. RNase P consists of three protein subunits: TRMT10C (MRPP1), SDR5C1 (MRPP2), and the endonuclease PRORP (MRPP3), which together mediate hydrolysis of the mt precursor tRNA (pre-tRNA) 5′ leader^1–5^. RNase Z consists of the same MRPP1/2 proteins as in RNase P, plus the ELAC2 endonuclease, which cleaves the pre-tRNA 3′ trailer (Fig. 1B)^4,5^. The MRPP1/2 complex is viewed as a “maturation platform”, positioning the pre-tRNA for reactions with the endonucleases^1,5,6^. Because mt-tRNAs are dispersed throughout the polycistronic transcript, they serve as punctuation marks that release the majority of individual mRNAs and rRNAs when processed. This “tRNA punctuation model” ensures the coordinated maturation of tRNAs, mRNAs, and rRNAs for OXPHOS protein expression within the mitochondrial matrix. The generally accepted stages in an mt-tRNA life cycle includes transcription, 5′ cleavage by RNase P, 3′ cleavage by RNase Z, post-transcriptional addition of the 3′-CCA tail by TRNT1, covalent modifications by “writer” enzymes, and aminoacylation by cognate tRNA synthetases (Fig. 1B)^5,6^.

**Figure 1.**
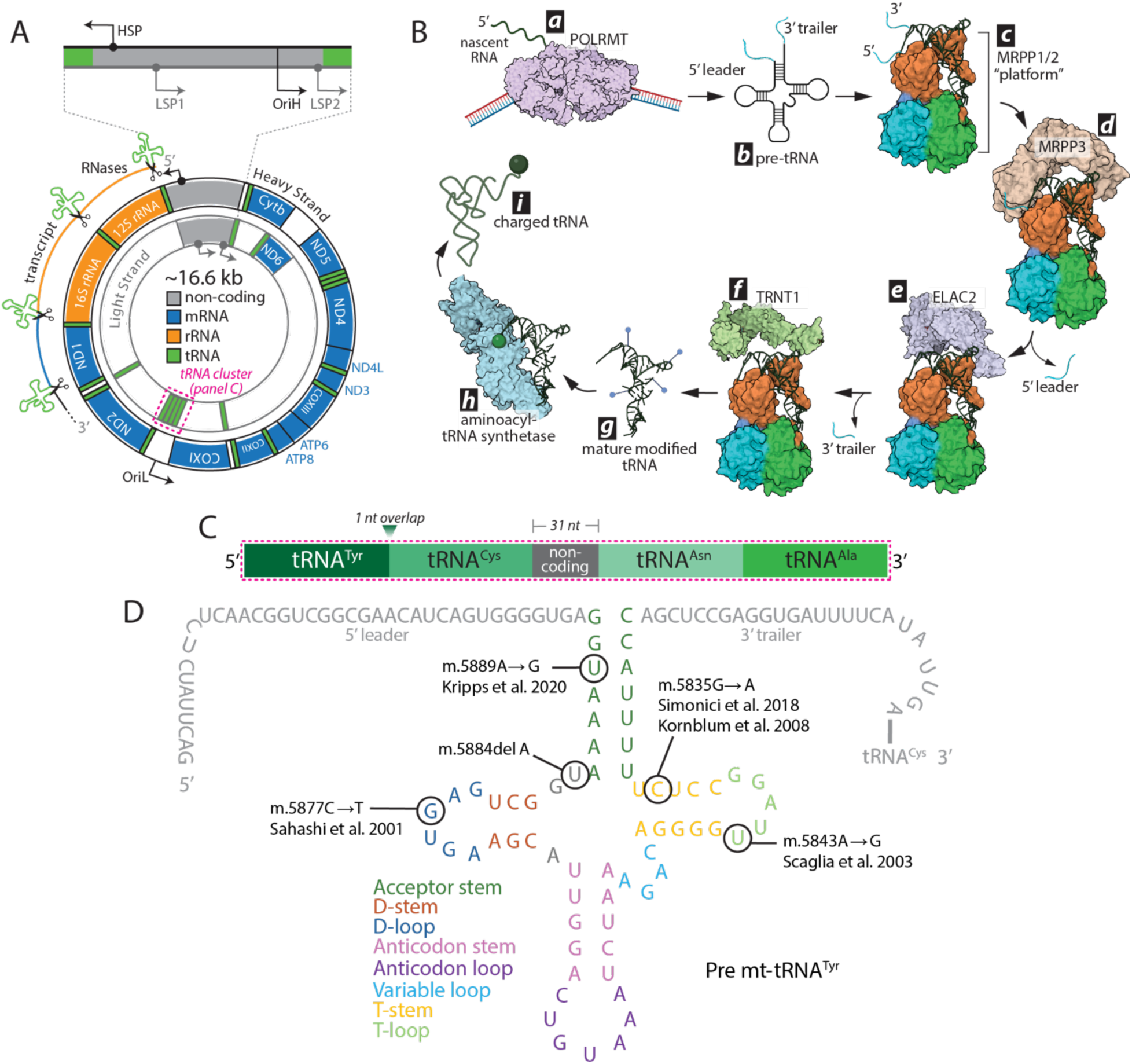
Mitochondrial tRNA life cycle and the mt-tRNA^Tyr^ variants explored in this study. (A) Map of human mitochondrial genome (mtDNA), with the tRNA^Tyr^ cluster highlighted in magenta. (B) Life cycle of mitochondrial tRNA starting with transcription by human mitochondrial RNA polymerase, POLRMT (*a*), nascent RNA folding into a precursor tRNA containing the 5′ leader and 3′ trailer (*b*), pre-tRNA association with by MRPP1/2 (*c*, PDB ID: 7ONU), 5′ leader processing by MRPP3 (*d*, PDB ID: 7ONU), 3′ trailer processing by ELAC2 (*e*, PDB: 8Z1G), 3′-CCA tail addition by TRNT1 (*f*, PDB ID: 4X4W), tRNA modification by “writer enzymes” (*g*), aminoacylation by the respective tRNA aminoacyl synthetase (*h*, PDB ID: 2PID), and mature mt-tRNA release (*i*). (C) Zoomed view of the light strand tRNA cluster with tRNA^Tyr^ as the pioneer tRNA transcribed. (D) Sequence of mitochondrial pre-tRNA^Tyr^ with the secondary structure elements and sites of the variants investigated in this work.

Defects in the above mt-tRNA processing steps have been linked to mitochondrial dysfunction and disease in model systems and patients^7,8^. For example, knockout of MRPP3 in mice reduces both 5′ and 3′ cleavage, impairing mitochondrial ribosome assembly and OXPHOS complex activity^9^. Loss of ELAC2 causes premature death in mice due to defective mt-tRNA processing and reduced OXPHOS function^10^. Collectively, such studies demonstrate that accurate 5′ and 3′ mt-tRNA cleavage is essential for mitochondrial function.

To date, 1,058 mtDNA variants have been reported, with ∼40% mapping to tRNA genes and 60% of those classified as pathogenic through a standardized scoring system developed by Clin Gen^11^. In this scoring framework, pathogenicity is determined by assigning weighted points to multiple lines of evidence, such as supported by multiple independent patient reports, evolutionary conservation of the affected nucleotide, heteroplasmy and segregation with disease, biochemical defects in OXPHOS complexes, functional and histochemical evidence of mitochondrial dysfunction, and the rarity or absence of the variant in large mtDNA databases^11^. Pathogenic mt-tRNA variants are associated with reduced OXPHOS complex production, and multiple biochemical studies have shown defects in 5′/3′ processing and aminoacylation due to mt-tRNA mutations^12–14^. No systematic studies have examined the impact of mt-tRNA single-point deletions in mt-DNA.

Although most human mt-tRNAs are separated by rRNA or mRNA genes, several regions of the mtDNA contain “tRNA clusters”, where multiple mt-tRNAs are transcribed consecutively (Fig. 1A). The sequential processing of individual tRNAs in tRNA clusters remains poorly understood, and it is unknown if mutations or deletions in the first transcribed mt-tRNA affect downstream mt-tRNA processing. One such cluster is present on the transcript from the light-strand promoter (LSP) and contains tRNA^Tyr^-tRNA^Cys^-tRNA^Asn^-tRNA^Ala^ (Fig. 1C). Because mt-tRNA^Tyr^ is positioned at the start of this cluster, its processing may act as a “gatekeeper”, such that changes in its cleavage efficiency alter downstream maturation events.

Here, we focused on the mt-tRNA^Tyr^ in its native cluster to establish a framework for *in vitro* investigation of disease-linked variants in cluster mt-tRNAs and how they may disrupt sequential processing. We investigated two disease-linked mt-tRNA^Tyr^ variants (m.A5889G^15,11^ and m.G5835A^16,17,11^) and a deletion variant (m.5884delA) located at or near the acceptor stem, as well as two “benign” variants (m.C5877T^11,18^ and m.A5843G^11,19^) within the D-loop and T-loop (Fig 1D). Benign variants provided a useful contrast, enabling direct comparison of folding and processing defects across variant classes.

Previous studies reported that m.C5877T and m.A5843G, although benign, reduce aminoacylation by tightening the T-loop tRNA^Tyr^ structure, though their effects on 5′/3′ cleavage remain unknown^13^. The m.G5835A variant dramatically reduced steady-state mt-tRNA^Tyr^ levels and aminoacylation (∼95%) in patient samples, but effects on processing were not assessed^16^. Thus, it remains unclear whether these pathogenic outcomes originate from early processing defects, misfolding, or downstream steps in tRNA maturation.

In this work, we analyzed the five *in vitro* transcribed pre-mt-tRNA^Tyr^ substrates for any effects of nucleotide sequence variation on the 5′ leader and 3′ trailer cleavage, MRPP1/2 binding, and processing of mt-tRNAs downstream of mt-tRNA^Tyr^ in the cluster. Our findings reveal the majority of examined variants compromise 5′ leader cleavage, which in turn reduces the yield of fully excised mt-tRNA^Tyr^ as well as the downstream mt-tRNA^Cys^ release. Some variants showed poor binding to MRPP1/2, explaining their reduced cleavage. Our work adds to the existing evidence in support of a hierarchical 5′-to-3′ mt-tRNA processing pathway in which MRPP3-mediated 5′ leader removal must occur before ELAC2 can act on the 3′ trailer, with MRPP1/2 playing an essential role by ensuring that this order is preserved. With these findings, we show that steps in early mt-tRNA processing are governed by 5′ processing and MRPP1/2-mediated pre-tRNA substrate orientation, with variations in the tRNA sequence compromising processing.

## MATERIALS AND METHODS

### Preparation of RNA substrates

Human mitochondrial DNA was purified from HEK293T cells, and region 5512-7585 (NCBI Reference Sequence: NC_012920.1) was cloned into a pUC19 vector. Variant mtDNA sequences containing either the deletion or mutations on mt-tRNA^Tyr^ within the light strand tRNA “cluster” were commercially synthesized as “E-blocks” by Integrated DNA Technologies and cloned into a pUC19 vector. All DNA sequences encoding the tRNA substrates are listed in Supplementary Table 1. Each plasmid was transformed into *E. coli* HB101 competent cells (Promega) and miniprepped using the ZymoPURE Plasmid Miniprep Kit (Zymo Research). These DNA constructs were used as PCR templates to prepare DNA for *in vitro* transcription. For the PCR, a forward primer was designed to be complementary to the 5′ leader sequence, and a reverse primer complementary to the 3′ trailer sequence. All primers are listed in Supplementary Table 2. The PCR products were gel-purified from a 1% agarose gel and extracted using the Monarch DNA gel extraction kit (New England Biolabs). The purified product was further PCR-amplified with a forward primer complementary to the 5′ leader sequence containing a T7 RNA Polymerase promoter overhang, and a reverse primer complementary to the 3′ trailer sequence. These final PCR-amplified products were cleaned up using the Monarch PCR and DNA clean up kit (New England Biolabs) and used for *in vitro* transcription by T7 RNA Polymerase.

All tRNA substrates used for biochemical analysis were transcribed using the HighScribe T7 Quick High Yield RNA Synthesis kit (New England Biolabs) in a 20 µL transcription reaction containing 0.20 µg of the DNA template, 3.3 mM NTP buffer mix, 2.5 mM DTT, 1 µL of T7 RNA Polymerase mix and incubated at 37 °C for 15 minutes. In the same tube, 70 µL of milliQ Nuclease Free water (ThermoFisher Scientific), 10 µL of Turbo DNase 10x buffer, and 2.5 µL Turbo DNase (ThermoFisher Scientific) were added to the reaction and incubated at 37 °C for an additional 20 minutes to degrade the transcription template DNA. The RNA was purified using the RNA Clean & Concentrator (Zymo Research). The RNA purity was verified on a 7% urea-PAGE gel run for 90 minutes at 100 V, then soaked in 1x Tris-Borate-EDTA buffer (ThermoFisher Scientific) containing 1x SYBR gold nucleic acid gel stain (ThermoFisher Scientific) and visualized on a Gel imaging system (BioRad).

### Protein cloning and expression

TRMT10C (MRPP1, UNIPROT: <u>Q7L0Y3</u>), SDR5C1 (MRPP2, UNIPROT: <u>Q99714</u>) and PRORP (MRPP3, UNIPROT: <u>O15091</u>) were synthesized and cloned into pUC57 vectors by Bio Basic with the predicted mitochondrial targeting signals (MTS) of MRPP1 (Δ1-39), MRPP2 (Δ1-11), and MRPP3 (Δ1-45) absent in each construct. Additionally, MRPP1 and MRPP3 contained an N-terminal 6x His-tag followed by a TEV protease cleavage site. MRPP1 and MRPP2 were cloned into a pETDuet-1 vector for co-expression. MRPP3 was also cloned into a pETDuet-1 vector (due to vector availability), with one of the promoters removed from the vector. ELAC2 (UNIPROT: <u>Q9BQ52</u>) was provided by colleagues Dr. Sean Reardon and Shannon Cole from the Mishanina Lab as a plasmid containing the MTS-lacking (Δ1-31) ELAC2 sequence with a SUMO protease cleavable N-terminal 6x His-tag in a pProEXHTB vector. All plasmids were transformed into *E. coli* DH5α cells for colony screening and plasmid propagation. Final expression plasmids were verified by Sanger sequencing before proceeding with protein expression. All plasmids listed in Supplementary Table 3.

MRPP1/2, MRPP3, and ELAC2 were each transformed into *E. coli* BL21(DE3) CodonPlus-RIPL cells (Promega) and selected for LB agar plates containing 100 μg/ml ampicillin and 34 μg/ml chloramphenicol overnight (12-16 h) at 37 °C. Starter liquid cultures were then prepared by inoculating 4 x 5 mL of LB broth containing 100 μg/ml ampicillin and 34 μg/ml chloramphenicol with one selected colony per 5 mL broth and growing overnight (12-16 h) at 37 °C with shaking. Each 5 mL starting overnight culture was inoculated into 1 L of LB broth containing 100 μg/ml ampicillin and shaken at 200 rpm at 37 °C until the OD_600_ reached 0.6 units (4-6 h). The temperature was then lowered to 16 °C, 0.5 mM (final) IPTG was added to the MRPP1/2 and MRPP3 culture flasks, and 0.15 mM (final) IPTG to the ELAC2 flask to induce expression of each respective protein, and the cultures were shaken overnight (16-20 h) at 16 °C.

### Protein purification

HEPES buffer was used for purification of MRPP1/2/3 (pH 7.5) and ELAC2 (pH 8.0) at 4 °C. Cell pellets were harvested and resuspended in the lysis buffer (50 mM HEPES, 500 mM NaC (for MRPP1/2/3) or 300 mM NaCl (for ELAC2) NaCl, 10 mM imidazole, 10% v/v glycerol, 1x Protease Inhibitor Cocktail (PIC: 31.2 µg/ml benzamidine, 0.5 µg/ml chymostatin, 0.5 µg/ml leupeptin, 0.1 µg/ml pepstatin, 1.0 µg/ml aprotinin, and 1.0 µg/ml antipain), 2 mM DTT, 1 mg/ml lysozyme (ThermoFisher Scientific). All protein purification steps were carried out at 4 °C, following the protocol from the Hillen lab as a starting point^2,5,20^. The resuspended cells were lysed by sonication for 20 minutes, 20 seconds off, 20 seconds on, 60% amplitude. Post-lysis, MRPP1/2 lysate was treated with an RNase cocktail (5 U/ml RNase A, 200 U/ml RNase T1, ThermoFisher Scientific), incubating at 4 °C and gently rocking for 2 hours. Each protein overexpression’s crude lysate was centrifuged at 30,187 x *g* for 30 minutes at 4 °C to pellet the cell debris. The supernatant was passed through a 0.45 µM filter before being applied to a 7.5 mL bed volume of Ni-NTA resin (Gold Bio) equilibrated with the respective lysis buffer and rocked at 4 °C (1 h for MRPP1/2, 30 minutes for MRPP3 and ELAC2). The resin was washed with 12 column volumes (CV) of high-salt wash buffer (50 mM HEPES, 1M NaCl, 20 mM imidazole, 10% glycerol, 1x PIC, and 2 mM DTT). Column-bound proteins were eluted with 2 CV of the appropriate elution buffer – for MRPP1/2: 50 mM HEPES, 300 mM NaCl, 400 mM imidazole, 10% glycerol, 1x PIC, and 1.7 mM DTT; for MRPP3: 42.5 mM HEPES, 425 mM NaCl, 317 mM imidazole, 8.5% glycerol, 1x PIC, and 1.7 mM DTT; for ELAC2: 42.5 mM HEPES, 255 mM NaCl, 317 mM imidazole, 8.5% glycerol, 1x PIC, and 1.7 mM DTT. The eluate was dialyzed overnight (16-20 h) against dialysis buffer (50mM HEPES, 200mM NaCl, 10% glycerol, 1x PIC, and 2mM DTT) in the presence of a recombinant His-tagged TEV protease (homemade; approximate molar ratio of 1:25 TEV:MRPP1/2/3) or recombinant ULP1 protease (homemade; approximate molar ratio of 1:500 ULP1:ELAC2). The dialyzed eluate was applied to the 7.5 mL bed volume of Ni-NTA resin equilibrated in each respective dialysis buffer and gently rocked for 20 minutes at 4 °C to allow binding of the cleaved His-tag and His-tagged protease. The untagged protein was eluted and washed with 1 CV of dialysis buffer, then applied to a 5 mL HiTrap heparin column (Cytiva) equilibrated with 3 CV heparin low salt wash buffer (50 mM HEPES, 200 mM NaCl, 10% glycerol, 1x PIC, 2 mM DTT) and washed with 5 CV of the heparin low salt wash buffer. Protein was then eluted with a gradient of 15-50% v/v heparin high salt elution buffer (50 mM HEPES, 1.5 M NaCl, 10% glycerol, 1x PIC, and 2 mM DTT). Fractions containing the protein of interest, identified by SDS-PAGE and Coomassie staining, were pooled and concentrated using Amicon 30K (for MRPP1/2/3) or 50K (for ELAC2) MWCO Centrifugal Filter Devices (ThermoFisher Scientific) and stored at −80 °C in storage buffer (50 mM HEPES, 150 mM NaCl, 25% glycerol, 1x PIC, and 2 mM DTT).

### RNA cleavage assays

All RNase P cleavage assays were carried out in 15 μl reactions containing RNA cleavage assay buffer (25 mM Tris-HCl pH 8.0 at 4 °C, 10 mM MgCl_2_, 0.1mg ml^−1^ BSA, 70 mM NaCl, 4 mM DTT, 20 µM *S*-adenosine methionine (ThermoFisher Scientific), 1 U/µl Murine RNase inhibitor (ThermoFisher Scientific) and 63 nM pre-tRNA^Tyr^ containing the native 38-nt 5′ leader and 24-nt 3′ trailer (same for the variants). The pre-tRNA^Tyr^ (NCBI Reference Sequence: NC_012920.1, GeneID:<u>4579</u>) was incubated with 650 nM MRPP1/2 at 37 °C for initial protein-RNA binding. After 10 minutes, each endonuclease was added either individually (MRPP3 for 5′ processing or ELAC2 without pre-MRPP1/2 for 3′ processing) or as a combination simultaneously to the reaction at 100 nM each protein final concentration and incubated for 30 minutes at 37°C. For extended substrate assays, such as pre-tRNA^Tyr^-tRNA^Cys^ and pre-tRNA^Tyr^-tRNA^Cys^-tRNA^Asn^, each substrate was pre-incubated with 650 nM MRPP1/2 for 10 minutes at 37 °C, reactions were initiated by the addition of the endonucleases (100 nM each final concentration) and incubated for 20 minutes. To capture additional tRNA released following initial processing, MRPP1/2 was subsequently added to a final concentration of 1.3 µM and reactions were incubated for an additional 10 minutes. MRPP3 and ELAC2 were then supplemented to a final concentration of 200 nM each and reactions were further incubated for 30 minutes. All reactions were quenched by the addition of 15 µL of 2x formamide-urea stop buffer (90 mM Tris-borate buffer, 8 M urea, 47.5 mM EDTA, 50% formamide, and 0.02% of each xylene cyanol and bromophenol blue) and stored at 4°C for up to a week without apparent RNA degradation. The full 30 µL reaction was carefully loaded onto 8 M urea 7% polyacrylamide (bis-acrylamide to acrylamide ratio 1:19) denaturing gels in 1x Tris-borate-EDTA buffer and run at 100 V for 120 minutes, 130 min (for the pre-tRNA^Tyr^-tRNA^Cys^-tRNA^Asn^ substrate), or 115 minutes (for the 5′ leaderless pre-tRNA^Tyr^). After electrophoresis, gels were stained with 1x SYBR Gold nucleic acid stain, visualized on a gel imaging system (BioRad), and quantified with ImageLab 6.1 software (BioRad). Each experiment was performed in 3-5 replicates, and the obtained data was analyzed using Prism version 10.6.1 (GraphPad) software.

### Native polyacrylamide gel electrophoresis (PAGE)

Each pre-tRNA^Tyr^ (2.4 pmol) was unfolded at 95°C for 3 minutes and placed on ice for 5 minutes before adding RNA folding buffer (25 mM HEPES pH 8.0, 7 0mM NaCl and either water or MgCl_2_ at the indicated final concentrations). After a 30 minute refold at room temperature, samples were mixed 1:1 with 2x native loading dye (1% bromophenol blue, 1% xylene cyanol, 100% glycerol) and run at 4°C on an 8.5% native polyacrylamide gel in 0.5x Tris-borate-EDTA buffer at 80 V, 400 mA for 140 minutes. Gels were then stained with 1x SYBR Green II (ThermoFisher Scientific) and visualized as previously described.

### Electrophoretic Mobility Shift Assays (EMSAs)

Each pre-tRNA^Tyr^ (55 nM final) was incubated with MRPP1/2 in varying concentrations in binding buffe r(50 mM Tris-HCl pH 8.0, 100 mM potassium acetate, 2.5 mM magnesium acetate, 1mM DTT, 1x murine RNase inhibitor) for 30 minutes at 37 °C. Glycerol was added to 5% v/v final concentration and each sample was loaded at 4 °C onto an 8.5% native polyacrylamide gel in 0.5x Tris-borate-EDTA buffer and run at 95 V, 400 mA for 106 minutes. Gels were then stained with 1x SYBR Green II (ThermoFisher Scientific) and then visualized and quantified as described.

Each experiment was performed in triplicate, and the obtained data was analyzed using Prism version 10.6.1 (GraphPad) software. Binding curves were fit and the K_d_ estimated using the specific binding with Hill slope equation:

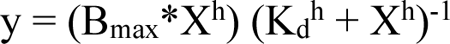

where B_max_ is the maximum specific binding, K_d_ is the mt-tRNA concentration needed to achieve a half-maximum binding at equilibrium, and h is the hill slope which >1.0 and generated a sigmoidal curve if there are multiple binding sites with positive cooperativity.

### UV melting experiments

Each pre-tRNA^Tyr^ (500 nM, wt or G5835A) was refolded in RNA refolding buffer using the method mentioned above and then RNA melting buffer (150 mM NaCl, 1 mM MgCl_2_, and 20 mM sodium cacodylate pH 7) was added. The pre-tRNAs melting was monitored on a Cary 100 UV Spectrophotometer (Agilent). Absorbance was measured at 260 nm as the temperature increased from 25 °C to 92 °C at a rate of 1 °C min^−1^ and T_m_ values were calculated from first-derivative plots (dA/dT vs T).

### Competition assay

Wild-type (wt) pre-tRNA^Tyr^ (40 nM final) was pre-mixed with increasing concentrations of pre-tRNA^Tyr^-tRNA^Cys^ in separate tubes prior to the addition of processing proteins, and reactions were carried out in a total volume of 15 µL as described above. All reactions were quenched by the addition of 15 µL of 2x formamide-urea stop buffer. After electrophoresis, gels were stained with 1x SYBR Gold nucleic acid stain and visualized and quantified as described above. Each experiment was performed in triplicate, and the obtained data was analyzed using Prism version 10.6.1 (GraphPad) software.

## RESULTS

### Pathogenically linked variants of the human mitochondrial tRNA^Tyr^ impair 5′ leader processing with minimal effect on 3′ trailer processing

To reconstitute human mitochondrial pre-tRNA processing *in vitro*, we recombinantly expressed and purified MRPP3 and ELAC2 individually and MRPP1/2 as a complex and *in vitro* transcribed the pre-tRNA^Tyr^ substrates (Fig. S1), each containing the native 38-nucleotide (nt) 5′ leader and the native 24-nt 3′ trailer. We analyzed the 5′-end processing by first forming an MRPP1/2-pre-tRNA complex and then adding the 5′ endonuclease MRPP3. MRPP3 is has been documented to require the presence of MRPP1/2 binding to carry out RNA cleavage^2,3,20,21^. Initial analysis of 5′ cleavage products revealed a significant reduction in cleavage activity in the A5889G, 5884delA, and G5835A pre-tRNA^Tyr^ variants, some reduced cleavage in A5843G, and wild-type (wt) pre-tRNA^Tyr^-like cleavage efficiency in the C5877T variant (Fig. 2A).

**Figure 2.**
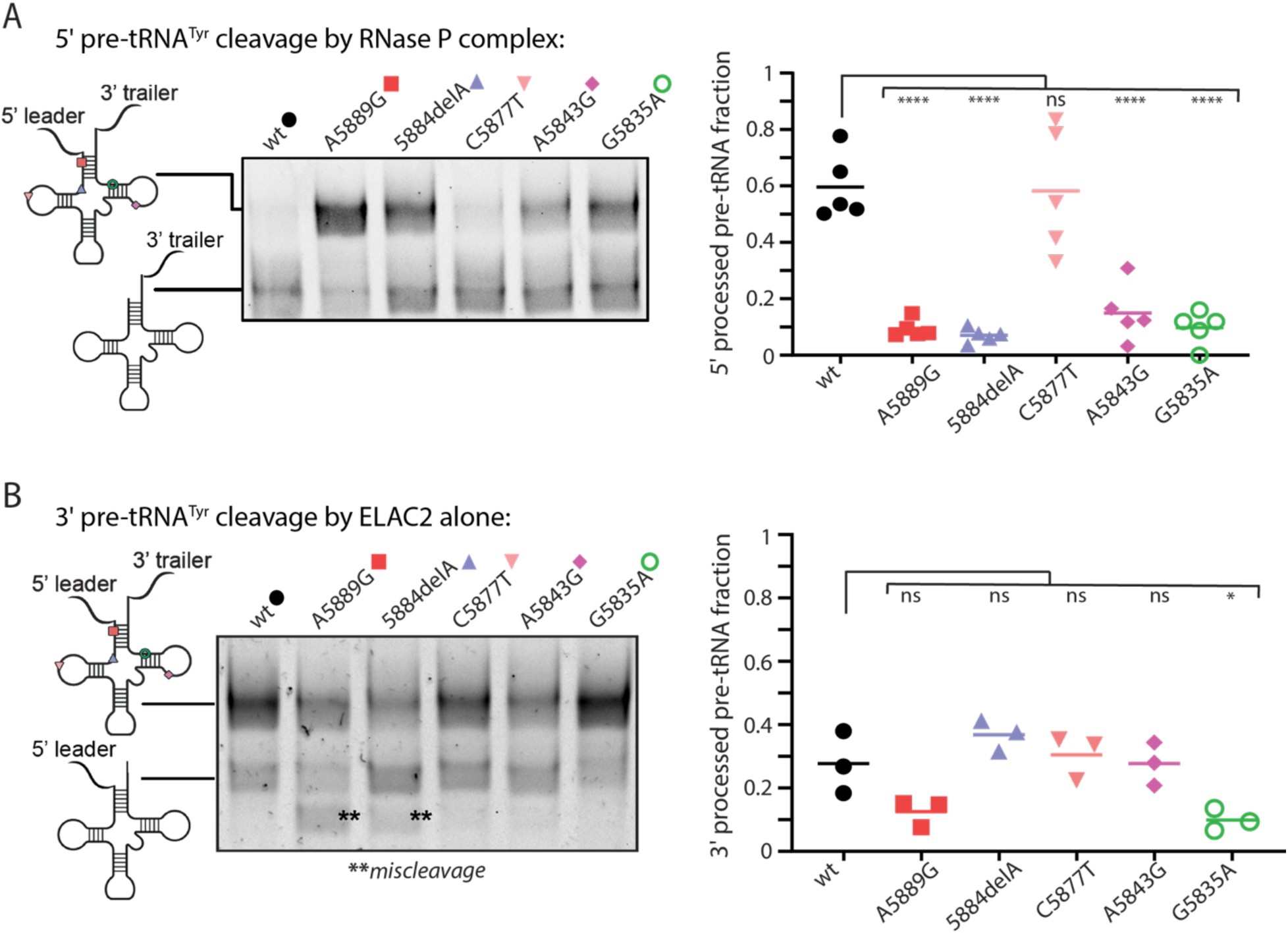
Sequence variants of mt-tRNA^Tyr^ impair 5′ leader processing by RNase P but not 3′ trailer processing by ELAC2 alone. (A) *In vitro* cleavage gel showing RNase P-catalyzed 5′ leader processing differences for the wild-type (wt) vs variant tRNA^Tyr^ precursors. The 5′ processed tRNA product fraction was quantified and plotted. (B) *In vitro* cleavage gel showing differences in 3′ trailer removal by ELAC2 alone on the wt vs variant tRNA^Tyr^. The 3′ processed tRNA fraction was quantified and plotted. A one-way ANOVA was used for statistical analysis (****adjusted *p*-value <0.0001, *adjusted *p*-value 0.0203).

To monitor the 3′ trailer hydrolysis, ELAC2 was added alone to the 5′ leader containing pre-tRNA substrates, as previous structural work by the Hillen lab showed that the 5′ leader of the MRPP1/2-bound pre-tRNA creates a steric clash with ELAC2, thereby preventing 3′ processing in the presence of the 5′ leader^5^. Analysis of 3′ cleavage patterns by ELAC2 in this context revealed generally similar processing efficiencies between wt pre-tRNA^Tyr^ and the variant substrates (Fig. 2B), although A5889G and 5884delA exhibited reproducible trends toward miscleavage, defined here as cleavage within sites of the pre-tRNA other than the cognate 3′ cleavage site (Figs. 2B and S2A).

To verify ELAC2 activity in the context of a fully assembled RNase Z complex, we carried out the 3′ hydrolysis reactions using 5′ leaderless pre-tRNA^Tyr^ substrates pre-bound to the MRPP1/2 platform. We observed similar 3′ cleavage trends across pre-tRNA^Tyr^ variants (Fig. S2B and S2C). Interestingly, the 5′ leaderless substrate cleaved with ELAC2 alone, i.e., in the absence of MRPP1/2, also resulted in reproducible miscleaved products (Fig. S2B and S2D).

### Full tRNA excision out of a polycistronic transcript is governed by the efficiency of 5′ leader processing

With the processing enzymes showing varying cleavage efficiencies and patterns across the pre-tRNA^Tyr^ variants, we next sought to assay cleavage when both endonucleases (MRPP3 and ELAC2) were present to replicate a physiological scenario. We assayed the one-pot 5′ and 3′ cleavage of both the wt pre-tRNA^Tyr^ and disease-associated variants by first combing the pre-tRNA substrate containing both the 5′ leader and 3′ trailer with MRPP1/2 and then adding MRPP3 and ELAC2 simultaneously. The resulting cleavage pattern closely resembled the 5′-only cleavage profile generated by RNase P (Fig. 3A, compare to Fig. 2A), with cleavage reduced in A5889G, 5884delA, A5843G, and G5835A and unchanged in the C5877T variant. When we tested different protein combinations with the wt pre-tRNA substrate containing both the 5′ leader and 3′ trailer, we observed that ELAC2 alone hydrolyzed some of the pre-tRNA (Fig. S3, lanes 3 and 6; consistent with Fig. 2B wt lane), whereas no cleavage of the 3′ trailer by ELAC2 occurred when the tRNA precursor was pre-bound by MRPP1/2 (Fig. S3, lane 7), in line with the inhibitory effect of MRPP1/2 on the ELAC2 enzymatic activity in the presence of the 5′ leader. In contrast, the presence of the complete RNase P complex capable of efficiently removing the 5′ leader (Fig. S3, lane 2) enabled ELAC2 to hydrolyze the 3′ trailer and yield a fully processed mt-tRNA^Tyr^ (Fig. S3, lanes 4 and 8; Fig. 2A wt lane). Together, these data support the ordered 5′ leader first-then 3′ trailer processing of a precursor mt-tRNA.

**Figure 3.**
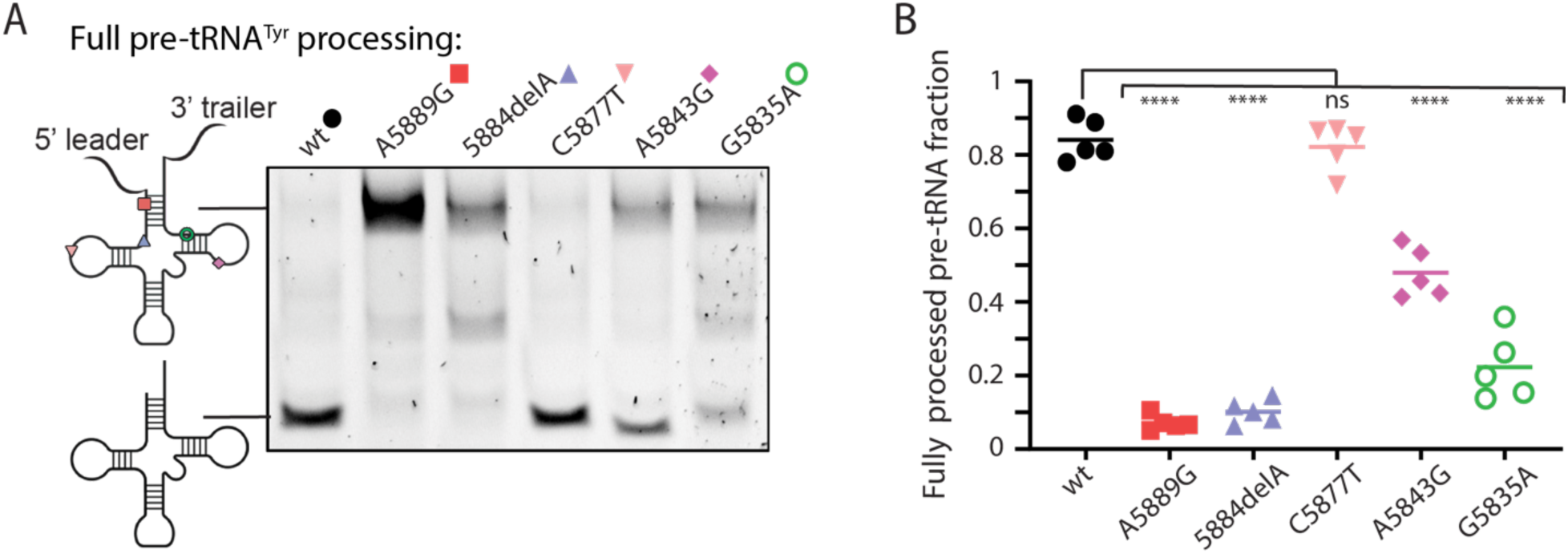
Sequence variants of mt-tRNA^Tyr^ compromise complete 5′ and 3′ cleavage by RNase P and RNase Z. (A) *In vitro* cleavage gel showing processing differences of the 5′ leader and 3′ trailer for the wt vs variant tRNA^Tyr^ precursors by RNase P and RNase Z. (B) The fraction of tRNA product without the 5′ leader and 3′ trailer was quantified and plotted. A one-way ANOVA was used for statistical analysis (****adjusted *p*-value <0.0001).

### Disease-associated sequence variations lower pre-tRNA^Tyr^ affinity for MRPP1/2

To gain insight into what step(s) in pre-tRNA processing the RNA sequence alterations impose their effect, we used electrophoretic mobility shift assay (EMSA) to qualitatively assess MRPP1/2-RNA binding. For consistency with the RNase-catalyzed cleavage experiments above, we used pre-tRNA^Tyr^ substrates containing the 5′ leader and 3′ trailer. Protein-bound RNA was quantified with increasing MRPP1/2 concentration, plotted and fitted to approximate K_d_ values using a Hill-slope model (Fig. 4, see Methods). The measured wt pre-tRNA^Tyr^ K_d_ (∼300 nM) was consistent with previously reported values determined by fluorescent anisotropy^5^. The C5877T and A5843G pre-tRNA^Tyr^ variants exhibited binding affinities akin to the wt pre-tRNA^Tyr^. The A5889G and 5884delA variants showed moderately increased K_d_ values indicative of weaker binding, whereas the G5835A mutant displayed a substantially higher K_d_ that exceeded the MRPP1/2 concentration range tested, suggesting significantly impaired binding.

**Figure 4.**
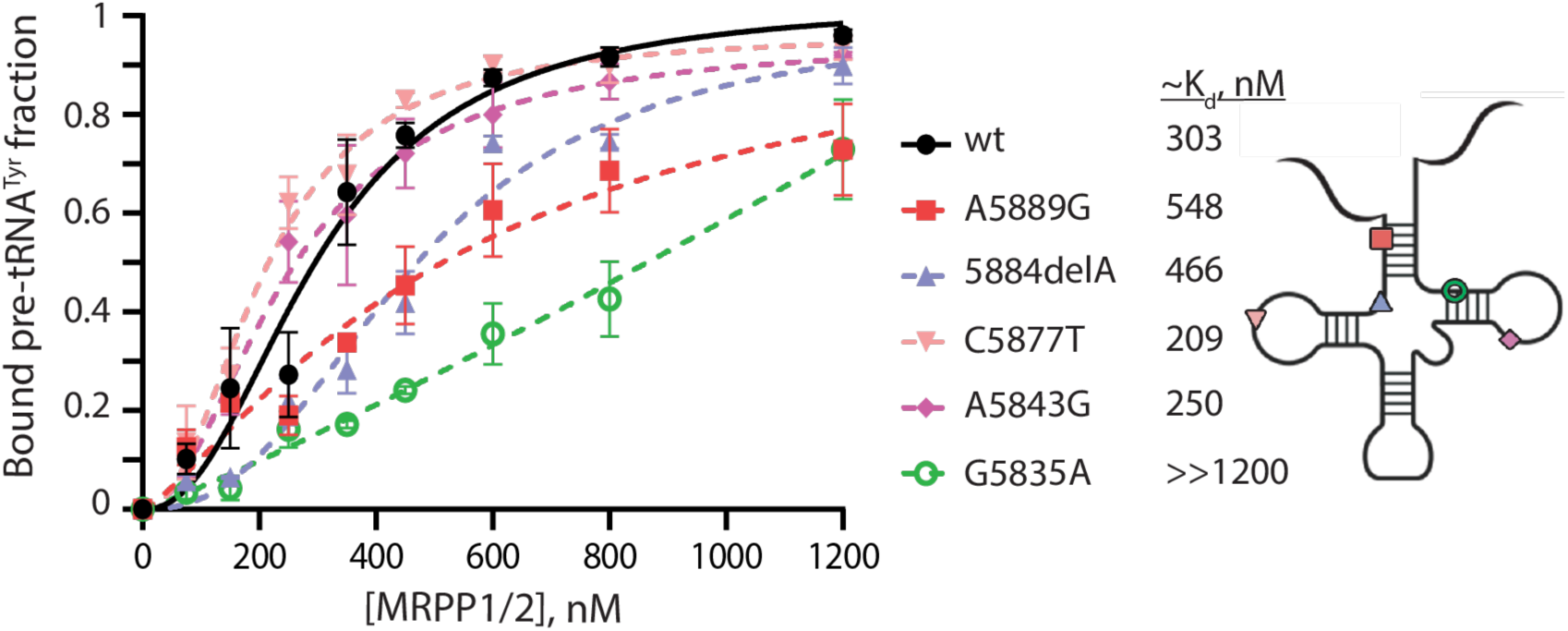
Mutations and a deletion in mt-tRNA^Tyr^ alter binding affinity for MRPP1/2. Quantitative analysis of pre-tRNAs bound by MRPP1/2. Each symbol represents the mean from three replicates with error bars showing the standard deviation. Protein-RNA dissociation constants (K_d_) was calculated in GraphPad Prism using a Hill-slope model (see Methods). Representative uncropped gel images are in Figure S7.

To assess whether variant pre-tRNAs assumed alternative conformations that may explain compromised association with MRPP1/2, we analyzed wt pre-tRNA^Tyr^ and variant pre-tRNAs by native PAGE. The wt pre-tRNA^Tyr^ displayed two predominant conformers, as evident from the two gel bands, and several variants showed a similar migration pattern (Fig. S4A). To further investigate the basis for the starkly weaker binding of the G5835A pre-tRNA^Tyr^ mutant to MRPP1/2, we performed thermal denaturation experiments comparing the stability of wt pre-tRNA^Tyr^ and G5835A pre-tRNA^Tyr^. At lower temperatures (∼35-45 °C), a slight shift in the first transition, associated with tertiary structural elements unfolding unfolding^21^, was observed. The second transition, corresponding to secondary structure melting, showed minimal differences between wt pre-tRNA^Tyr^ and the G5835A mutant in first-derivative plots (Fig. S4B), ∼60 °C and 62 °C respectively, consistent with previous UV melting data showing reduced peak amplitude of the derivative maxima in mutant pre-tRNAs^21^. This pattern persisted at higher temperatures (∼70-90 °C). We next examined whether Mg^2+^ concentration in solution influenced pre-tRNA conformational states, given that Mg^2+^ plays a critical role in stabilizing tertiary architecture in cytosolic tRNAs via a site-specific coordination^22^. Native gel analysis of the wt pre-tRNA^Tyr^ and the G5835A mutant performed at 0-10 mM Mg^2+^ showed similar migration patterns, with two dominant conformers observed once again across all divalent metal ion conditions (Fig. S4C).

An open question in the field is when in the pre-tRNA life cycle MRPP1/2 releases the tRNA with current models proposing that MRPP1/2 remains bound to the tRNA for 5′ leader and 3′ trailer hydrolysis, as well as 3′ CCA tail addition^5,6^. Because the A5843G pre-tRNA^Tyr^ variant bound MRPP1/2 with a similar relative affinity to the wt pre-tRNA^Tyr^ (Fig. 4), yet showed decreased cleavage (Figs. 2 and 3), we tested whether the A5843G mutant tRNA precursor could act as an MRPP1/2 “sponge” to sequester this protein complex and consequently impair wt pre-tRNA^Tyr^ processing. To model increasing mtDNA mutant loads (heteroplasmy) under physiologically relevant conditions, we performed 5′ cleavage assays using a substrate mixture of a constant amount of the wt pre-tRNA^Tyr^ and increasing concentration of the A5843G mutant. Processing of the 5′ leader remained unaffected by the presence of the A5834G pre-tRNA^Tyr^, even when the mutant concentration exceeded that of the wt pre-tRNA^Tyr^ (Fig. S4D and S4E).

### Release of mature mt-tRNA^Cys^ from the tRNA cluster is dependent on its 5′-proximal mt-tRNA^Tyr^ processing

Because tRNA^Tyr^ and tRNA^Cys^ are transcribed in tandem on the light strand of mtDNA and overlap by one nucleotide (Fig. 1C), we wondered whether the tRNA^Cys^ processing depends on the successful processing of the preceding tRNA^Tyr^. To address this, we *in vitro* transcribed extended pre-tRNA substrates consisting of the tRNA^Tyr^ sequence with its native 38-nt 5′ leader, the full tRNA^Cys^ and its native 48-nt 3′ trailer, and performed the same 5′ and 3′ cleavage assays as with the stand-alone pre-tRNA^Tyr^ (Fig. 3). The tRNA^Tyr^ variants that showed reduced cleavage on isolated pre-tRNA^Tyr^ substrates also displayed substantially decreased release of mature tRNA^Tyr^ and tRNA^Cys^ in the extended constructs (Fig. 5).

**Figure 5.**
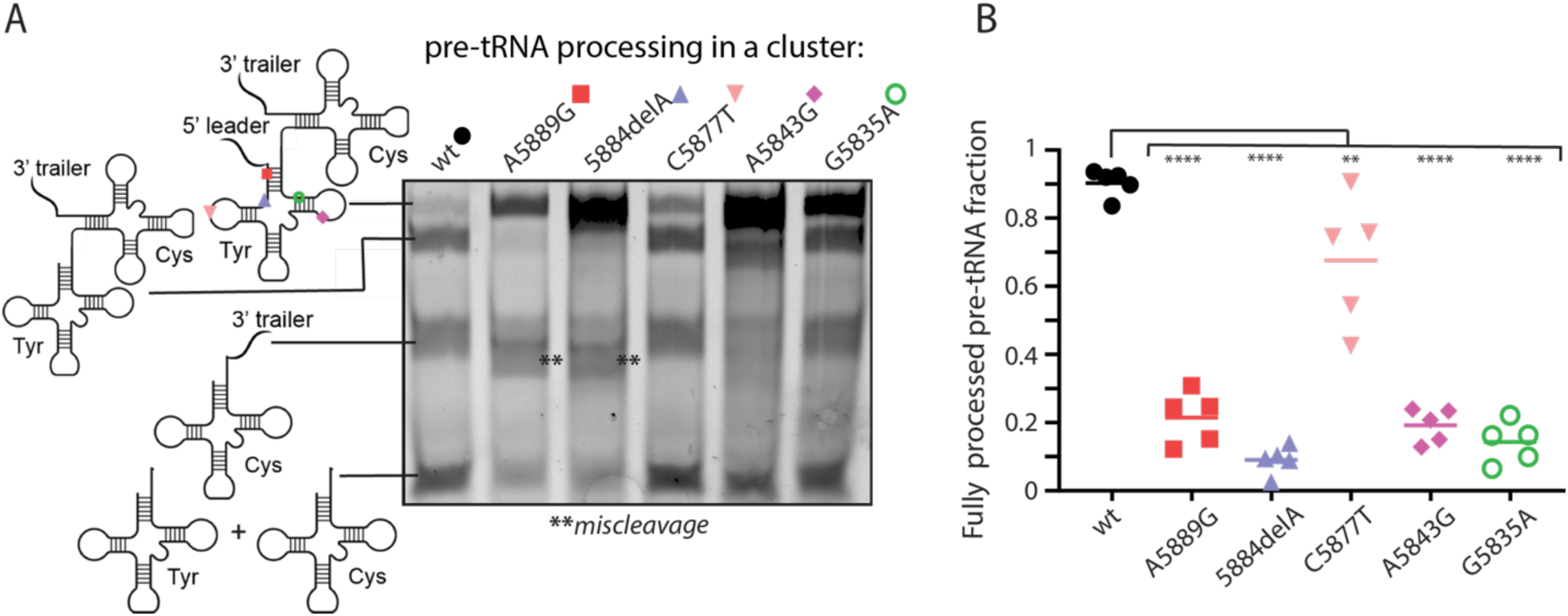
Processing of mt-tRNA^Cys^ is influenced by the sequence variation within the preceding tRNA^Tyr^. (A) *In vitro* cleavage gel showing differences in processing of the 5′ leader and 3′ trailer on the wt vs tRNA^Tyr^ precursors containing the tRNA^Tyr^-tRNA^Cys^ extended sequence. ** denotes miscleaved products within the tRNA^Tyr^. (B) The fraction of tRNA product without the 5′ leader and 3′ trailer was quantified and plotted. A one-way ANOVA was used for statistical analysis (****adjusted *p*-value <0.0001, ** adjusted *p*-value 0.0031).

We next examined the processing of the tRNA^Tyr^-tRNA^Cys^-tRNA^Asn^ cluster, as these tRNAs belong to the same light-strand transcript. A 31-nt noncoding region separates tRNA^Cys^ and tRNA^Asn^ (Fig. 1C) the tRNA^Asn^ processing, thus may not rely on the tRNA^Cys^ release, which itself in turn depends on its 5′-adjacent tRNA^Tyr^ processing. Previous work from the Guan lab using mitochondrial cybrids demonstrated that a tRNA^Cys^ mutation decreased mature tRNA^Cys^ as well as tRNA^Tyr^, while tRNA^Asn^ levels remained unchanged^23^. Consistent with these observations, tRNA^Asn^ was efficiently excised from the extended tRNA^Tyr^-tRNA^Cys^-tRNA^Asn^ transcript in our assays (Fig. S5).

## DISCUSSION

### Human mitochondrial pre-tRNA sequence variants and their effect on the early stages of the tRNA life cycle

Recent advances in visualizing high-resolution 3D structures of the mitochondrial RNase P and RNase Z complexes engaged with a tRNA precursor have beautifully illuminated how these enzymes recognize and cleave pre-tRNA substrates^5,6^. The impact of nucleotide variation within the pre-tRNAs on these processing steps, however, remains poorly understood. 60% of mutations, deletions and insertions in mt-tRNAs are associated with mitochondrial dysfunction, yet a systematic understanding of how these sequence changes influence the 5′ and 3′ processing of the tRNA is lacking. Earlier studies demonstrated that some individual variants perturb either 5′ or 3′ cleavage^21,23,24^, but little is known about dual-end processing, the effects of deletions, or how processing of one tRNA influences excision of adjacent tRNAs within the same transcript. Here, we expand upon these prior findings by defining how specific variations within human mt-tRNA^Tyr^ alter enzyme recognition, cleavage efficiency, and downstream maturation within the multi-tRNA transcriptional unit.

Some human mt-tRNAs are transcribed as a cluster without separation by rRNA or mRNA. In the light-strand cluster containing tRNA^Tyr^-tRNA^Cys^-tRNA^Asn^-tRNA^Ala^, tRNA^Tyr^ processing may regulate downstream maturation of the sequential mt-tRNA release (Fig. 1C). Several disease-associated sequence variants of mt-tRNA^Tyr^ have been linked to disorders including myopathy, neurological deterioration, and chronic progressive external ophthalmoplegia^15–19^ (Fig. 1D). However, the mechanistic connection between early tRNA transcript processing and these disease phenotypes has remained unclear. Our analysis provides evidence that the impact of these variants depends on their location within the mt-tRNA and proximity to enzyme interaction surfaces, offering a framework for understanding how early processing alterations may contribute to mitochondrial disease. Here we show how the specific nucleotide substitutions – along with a single-nucleotide deletion – within the mt-tRNA^Tyr^ negatively influence its 5′ leader processing by RNase P, with minimal effects on the 3′ trailer processing by RNase Z. The A5889G and 5884del, both located in or proximal to the tRNA acceptor stem, produced the largest reduction in 5′ cleavage, likely due to their proximity to the MRPP3 catalytic site (Figs. 2 and 6). A previous study found that the A4269G mutation in the acceptor stem of human mt-tRNA^Ile^ decreased 5′ processing by disrupting the base-pairing at the base of the acceptor stem^22^. Similarly, the A5889G substitution may disrupt base-pairing in the pre-tRNA^Tyr^, influencing how MRPP3 engages with the substrate. RNAfold simulations show that the A5889G centroid structure – the average structure from all potential structures – disrupts acceptor-stem base pairs and widens the overall tRNA architecture, in contrast to the cloverleaf structure of the wt pre-tRNA^Tyr^ (Figs. S6A and S6B). Consistent with the secondary structure predictions, the A5889G variant displays nearly a two-fold increase in its K_d_ for MRPP1/2 relative to the wt pre-tRNA^Tyr^ (Fig. 4), indicating weaker association with the MRPP1/2 processing platform. The Koutmos lab observed similar compromised MRPP1/2 binding for their pre-tRNA^Ile^ and pre-tRNA^Leu^ containing mutations in the acceptor stem^21^.

**Figure 6.**
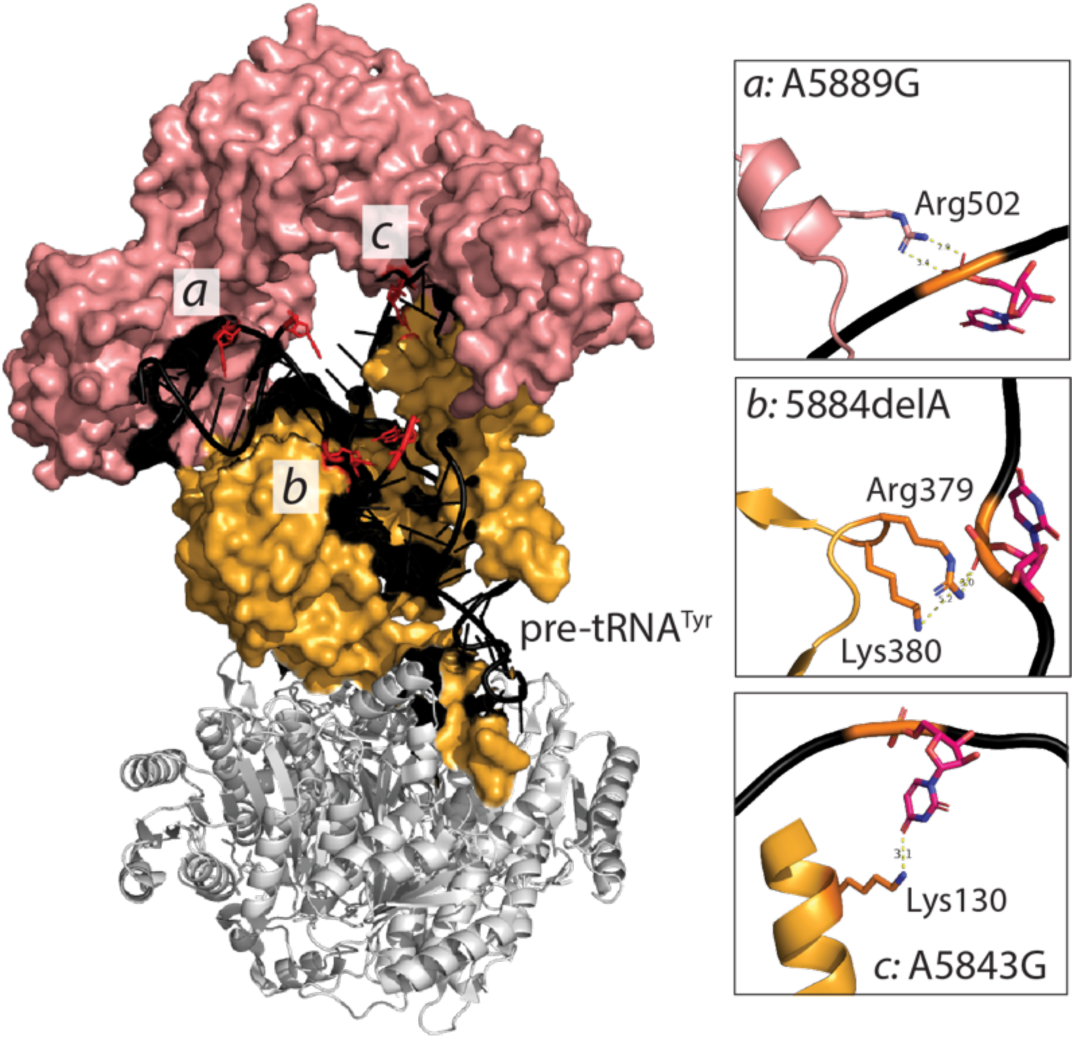
Interactions of the tRNA^Tyr^ substrate with RNase P. 3D structure and interactions (black patches) of mitochondrial pre-tRNA^Tyr^ within the mitochondrial RNase P complex (MRPP1 gold; MRPP2 gray; MRPP3 endonuclease salmon pink. PDB ID: 7ONU). Protein-RNA interactions for the representative variants with impaired 5′ leader cleavage are shown in the insets. Majority of protein interactions are with the phosphate-sugar backbone, except for m.A5843 (uracil in the RNA) where MRPP1 lysine and the carbonyl of the uracil ring form base-specific hydrogen bonding interactions (inset *c*).

The 5884delA deletion shortens the distance between the acceptor stem and the D-stem, potentially creating a more constricted substrate that hinders access of MRPP3 to the 5′ leader (Fig. S6C). MRPP1/2 binding to this variant is also impaired, though to a lesser extent than in A5889G (Fig. 4). This may be because the anticodon region, which MRPP1/2 engages with most extensively, remains largely intact in 5884delA, whereas A5889G disrupts interactions more globally (Figs. 6 and S6C).

With the C5877T mutation located in the D-loop, the cleavage efficiency of its 5′ leader and binding to MRPP1/2 closely matched wt pre-tRNA^Tyr^ (Figs. 2, 4, S7A and S7D). Consistent with these observations, the predicted secondary structure shows only minimal changes due to this substitution (Fig. S6A and S6D). A previous study using chemical and enzymatic probing, however, showed that this nucleotide substitution contributes to the structural destabilization of the tRNA D-arm and a 40-fold reduction in aminoacylation^13^. This residue may participate in transient tertiary interactions, potentially corresponding to the weaker, less stable conformer detected in our native gels and does not have direct interactions with the MRPP1/2 or endonuclease surface residues (Figs. S4A, S6, and S7D).

The A5843G mutation in the T-loop was also characterized by Bonnefond et al. for aminoacylation and showed only a modest (2-fold) reduction in the tRNA^Tyr^ charging with an amino acid^13^. The predicted secondary structure closely resembles that of the wt pre-tRNA^Tyr^, which is consistent with its near-wild-type pre-tRNA^Tyr^ binding to MRPP1/2 (Figs. 4 and S7E). Nevertheless, the 5′ cleavage efficiency of this mutant by MRPP3 was lower than for the wt pre-tRNA^Tyr^, possibly due to altered interactions between the mutated T-loop region and non-catalytic surfaces of MRPP1/2 or MRPP3. The T-loop is positioned closer to these enzyme interfaces than the D-loop, having interactions with both MRPP1/2 and each endonuclease, which could explain the drop in 5′ processing (Fig. 6). Our competition assays showed that the A5843G pre-tRNA^Tyr^ substrate does not inhibit 5′ cleavage of the wt pre-tRNA^Tyr^ even at elevated concentrations of the mutant competitor (Figs. S4D and S4E), suggesting that this variant would likely impact physiology only at high mutant mt-DNA levels, consistent with the >70% heteroplasmy reported in patients^20^. Notably, this variant is not classified as pathogenic according to criteria such as conservation, heteroplasmy, segregation, biochemical defects, and cybrid evidence^11^.

The final mutation, G5835A, located in the T-stem, affected both the 3′ trailer processing and 5′ leader cleavage – unlike all other variants, which primarily compromised 5′ leader processing (Fig. 2). This variant showed severe defects in binding to MRPP1/2 (Figs. 4 and S7F), consistent with the near absence of cleavage. Patient cell lines harboring this mutation lacked mature and aminoacylated mt-tRNA^Tyr^, in line with our biochemical *in vitro* findings where this variant fails to be processed out of the polycistronic transcript^16^. RNAfold predicts a large unpaired region following the D-loop, which may destabilize or break hydrogen-bonding interactions (Fig. S6F). Thermal denaturation analyses showed that although the first melting transition of the G5835A variant (∼60 °C) was similar to the wt pre-tRNA^Tyr^, the maximum of the first-derivative curve was markedly lower, suggesting an altered structural ensemble with reduced stabilizing interactions (Fig. S4B). At higher temperatures, resolution of transitions became more difficult, likely due to the presence of two structural conformers, visible on native gels that unfold differently (Fig. S4C). The ∼2 °C difference in melting temperature for the first curve resembles observations from a T-stem mt-tRNA^Cys^ variant and parallels behavior seen in mt-tRNA^Ile^ and mt-tRNA^Ser^ mutants, where melting temperatures remain similar but the derivative maxima shift^21,23^. Interestingly, the nucleotides do not appear to have apparent interactions with MRPP1/2, MRPP3, or ELAC2 (Fig. 6). Definitive determination of the structural differences imposed by the G5835A tRNA^Tyr^ mutation will require future RNA structure-probing experiments.

### The role of MRPP1/2 in the sequential 5**′**-to-3**′** mitochondrial pre-tRNA processing pathway

Our data suggest a model where full maturation of mt-tRNAs depends primarily on successful 5′ processing. This idea is consistent with observations from a study that knocked out MRPP3 in mice, which resulted in reduced 5′ and 3′ mt-tRNA cleavage and led to defects in the mitochondrial ribosome assembly and OXPHOS activity^9^. Structural studies from the Hillen lab also indicate that the 5′ leader creates a steric clash within the ELAC2 active site when MRPP1/2 is bound to the pre-tRNA, implying that RNase P-mediated 5′ cleavage is required *before* ELAC2 can properly engage with the substrate^5^. This does not mean that ELAC2 cannot function independently as we also observed 3′ cleavage when ELAC2 acted alone, without the MRPP1/2 maturation platform (Fig. 2B). This aligns with findings from the Churchman lab showing that some mt-RNAs possess 3′ adenylated tails even when 5′ leaders remain unprocessed^14^. Nonetheless, we observed MRPP1/2 inhibition of ELAC2 catalysis when the 5′ leader was present (Fig. S3B). Together, these studies indicate that ELAC2 has independent catalytic activity, but MRPP1/2 dictates the correct order of cleavage events.

We additionally observed miscleavage by ELAC2 in wt pre-tRNA^Tyr^ when MRPP1/2 was absent (Fig. S2A, S2B, and S2D). ELAC2 is known to have broad substrate tolerance, but its accuracy on mt-RNA substrates is less understood – particularly because many mt-tRNAs cannot form the canonical tertiary “elbow” architectures due to lacking key nucleotides in the D-loop and T-loop^14^. Prior studies suggested that mitochondrial RNase P components help orient non-canonical substrates for cleavage, and our data support this idea as miscleavage was detected when ELAC2 acted alone but disappeared when MRPP1/2 was included or when the 5′ leader was removed (Fig. 2B, Fig. S2A, Fig. S3B)^2,3,5,6,25^. Moreover, the miscleavage detected on the wt pre-tRNA^Tyr^ was not observed in full 5′ leader and 3′ trailer removal assays, further reinforcing the sequential model of mt-tRNA maturation (Fig. S2C).

### Processing of downstream mt-tRNA^Cys^ affected by mt-tRNA^Tyr^ variants

Our extended-substrate processing assays reveal that mt-tRNA^Cys^ processing depends strongly on the successful excision of the upstream mt-tRNA^Tyr^ (Fig. 5). This is perhaps expected as tRNA^Tyr^ and tRNA^Cys^ overlap by one nucleotide, meaning release of tRNA^Cys^ requires accurate and timely tRNA^Tyr^ 3′ cleavage. Such a model is supported by a prior study which showed using mitochondrial cybrids that the m.C5783T mutation on the T-stem of tRNA^Cys^ reduced mature levels of both tRNA^Cys^ and tRNA^Tyr^, likely as a direct consequence of impaired tRNA^Cys^ 3′ processing^23^.

To extend the tRNA cluster further, we examined tRNA^Tyr^-tRNA^Cys^-tRNA^Asn^ processing. Variants that impaired cleavage of the tRNA^Tyr^-tRNA^Cys^ portion did not significantly impact release of tRNA^Asn^. This is consistent with the presence of a 31-nt non-coding region between tRNA^Cys^ and tRNA^Asn^, allowing RNase P to cleave the 5′ end of tRNA^Asn^ independently of tRNA^Cys^release. Prior studies showed that tRNA^Asn^ can be processed by RNase P or ELAC2 alone and that tRNA^Ala^ – which follows tRNA^Asn^ – can be cleaved by MRPP3 independently of MRPP1/2^5,26^. Notably, tRNA^Asn^ and tRNA^Ala^ are among the few mt-tRNAs predicted to adopt canonical cytosolic-like tertiary structures, as they retain the nucleotide features required for the classical D-T loop tertiary fold^5^. Meng and Jia similarly observed normal tRNA^Asn^ abundance in m.C5783T tRNA^Cys^ mutation-containing cybrids^23^. Together, the published results and our current work suggest that downstream RNA processing defects are likely only when a tRNA relies directly on the release of its upstream neighbor. If no such dependency exists (such as having a non-coding spacer), processing proceeds normally as long as the relevant enzymes can recognize the pre-tRNA substrate.

## CONCLUSIONS

We found that mutations and a single-nucleotide deletion within the precursor tRNA can impair mitochondrial tRNA processing, often by altering local structure in ways that reduce MRPP3, ELAC2, or MRPP1/2 accessibility and binding. These defects likely arise from local changes in base-pairing or overall pre-tRNA conformation, each of which can disrupt recognition by the RNA processing machinery. Our work also supports a key role for MRPP1/2 in maintaining RNA processing order in human mitochondria. Variants in the pre-tRNA^Tyr^ acceptor stem produced some of the strongest defects, with further investigation needed to fully distinguish contributions from binding vs catalytic positioning. Moreover, the question of whether multiple-nucleotide deletions disrupt processing to the same extent as single-nucleotide deletions remains to be explored. Finally, our data shows that mutations in and upstream mt-tRNA can affect processing of downstream tRNAs in a tRNA cluster, but only when the downstream mt-tRNA relies directly on accurate 3′ processing of the upstream tRNA, e.g. due to sequence overlap. This principle may explain why some disease-associate variants disrupt only a subset of tRNAs within a polycistronic transcript, helping to connect molecular processing defects with patient phenotypes.

## ASSOCIATED CONTENT

### Supporting Information

Purification gels for preparation of the recombinant human RNase P and RNase Z and tRNA substrates; 3′ miscleavage by ELAC2 on full-length and leaderless pre-tRNA^Tyr^ gel image and plotted graphs; wild-type pre-tRNA^Tyr^ cleavage in the presence different protein combinations gel images; variant effects on tRNA conformation and competitive inhibition of 5′ leader removal with native PAGE gel images, UV-vis RNA melting curve, and plotted processed 5′ wt pre-tRNA^Tyr^; release of tRNA^Asn^ from the tRNA^Tyr^-tRNA^Cys^-tRNA^Asn^ cluster gel image and band quantitation plotted; RNAfold centroid structure models of the wild-type and variant tRNA^Tyr^; representative EMSA gels for wt and variant pre-tRNA^Tyr^; full gel images for biological replicates for individual 5′ and 3′ processed tRNA^Tyr^; full gel images for biological replicates for dual 5′ and 3′ processed tRNA^Tyr^; full gel images for biological replicates for dual 5′ and 3′ processed tRNA^Tyr^-tRNA^Cys^ (<u>PDF</u>)

Table of DNA templates for tRNA *in vitro* transcription; Table of primers; Table of plasmids (<u>XLSX</u>)

### Accession codes

TRMT10C (MRPP1, UNIPROT: <u>Q7L0Y3</u>)

SDR5C1 (MRPP2, UNIPROT: <u>Q99714</u>)

PRORP (MRPP3, UNIPROT: <u>O15091</u>

ELAC2 (UNIPROT: <u>Q9BQ52</u>)

## AUTHOR INFORMATION

### Author Contributions

The manuscript was written through contributions of all authors. All authors have given approval to the final version of the manuscript. J.H.M., conceptualization; J.H.M., methodology; J.H.M. and A.O., investigation; J.H.M., formal analysis; J.H.M., visualization; J.H.M., writing-initial draft; J.H.M., and T.V.M, writing-review and editing; T.V.M, supervision; T.V.M., funding acquisition.

### Notes

The authors declare no competing financial interests.

## ACKNOWLEDGMENTS

We would like to thank Dr. Susan Ackerman for her valuable insights in project development and direction. We would also like to thank Dr. Phil Ordoukhanian (Scripps Research Institute) for his training and guidance on the use of the Cary 100 Spectrophotometer. We would like to thank Dr. Elizabeth Komives for her encouragement. We would like to thank and acknowledge Linh Tran and Aaron Bagaoisan for their support and early-stage assistance for this project. We are grateful to Jonathan Eger for assistance with figure preparation. We greatly appreciate Yuntong Ou and Shannon Cole for the assistance with various components such as gel staining and providing protocols that aided us in this work. We are also greatly appreciative of Dr. Mukesh Mahajan, Dr. An Hsieh, Dr. Sean Reardon, and Dr. Ravish Sharma for their protocol refinement suggestion and intellectual advice and feedback. Thank you to the rest of the Mishanina research group for their feedback and encouragement.

## ABBREVIATIONS

OXPHOS: oxidative phosphorylation
mtDNA: mitochondrial DNA
ATP: adenosine triphosphatePOLRMT
POLRMT: human mitochondrial RNA polymerase
mRNA: messenger RNA
rRNA: ribosomal RNA
tRNA: transfer RNA
MT: mitochondrial
MRPP1: Mitochondrial RNase P protein 1 (TRMT10C, tRNA methyltransferase 10C)
MRPP2: Mitochondrial RNase P protein 2 (SDR5C1, Succinate dehydrogenase complex assembly factor 1)
MRPP3: Mitochondrial RNase P protein 3 (PRORP, Protein-only RNase P)
ELAC2: ElaC ribonuclease Z 2
RNase P: MRPP1/2/3
RNase Z: MRPP1/2 & ELAC2
MTS: mitochondrial targeting signal
PIC: protease inhibitor cocktail

