## Supplemental Information for "Pathogenic human mitochondrial tRNA variants impair RNA processing by compromising 5′ leader removal"

Contents:

Figures S1-S11

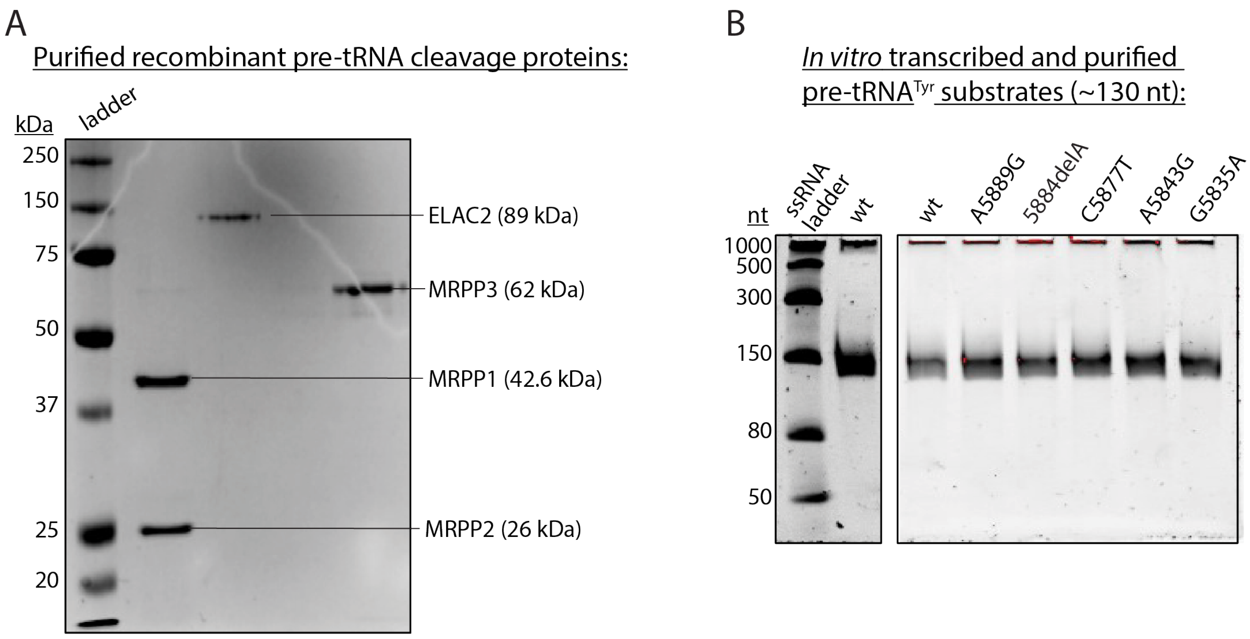


**Figure S1 | Preparation of the recombinant human RNase P and RNase Z and tRNA substrates | (A)** 10% SDS-PAGE showing purified protein components of the human mitochondrial RNase P and RNase Z, stained with Coomassie. **(B)** 7% urea-polyacrylamide gel showing purified pre-tRNA$^{Tyr}$ substrates for cleavage, stained with 1x SYBR Gold.

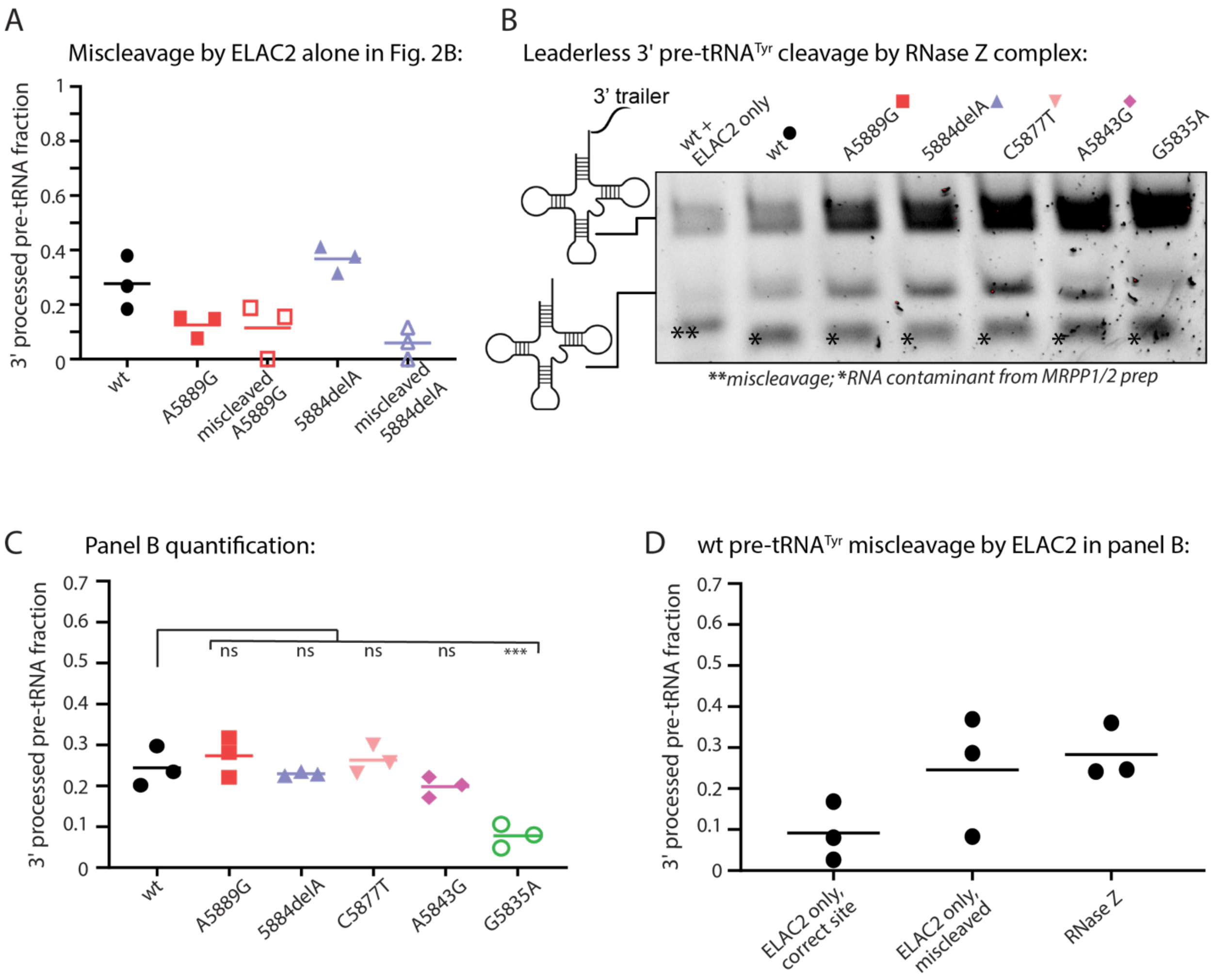


**Figure S2 | 3′ miscleavage by ELAC2 on full-length and leaderless pre-tRNA$^{Tyr}$ | (A)** Graph showing the 3′ processed fraction of the wt vs the A5889G and 5884delA tRNA$^{Tyr}$ variants with their respective miscleaved products alongside the correctly 3′ processed products. Data from the Fig. 2B gel, pixel intensities quantified using ImageLab (BioRad) and plotted using GraphPad prism. **(B)** *In vitro* cleavage assay showing processing of the 3′ trailer on the pre-tRNA$^{Tyr}$ lacking 5′ leader. RNA resolved on a 7% urea-polyacrylamide gel, stained with 1x SYBR Gold, representative of three replicates. **3′ trailer miscleavage of the substrate, *RNA contaminant that we could not get rid of during the recombinant preparation of MRPP1/2. **(C)** Graph showing the fraction of 3′ processed products of the wt pre-tRNA$^{Tyr}$ compared to the variants. Measured from Fig. S2B gel, quantified with ImageLab (BioRad) and plotted using GraphPad prism. **(D)** Graph showing the fraction of 3′ processed wt pre-tRNA$^{Tyr}$ cleaved correctly by ELAC2, miscleaved by ELAC2, and cleaved correctly by RNase Z complex (ELAC2 plus MRPP1/2). Measured from Fig. S2B gel, quantified with ImageLab (Biorad) and plotted using GraphPad prism.

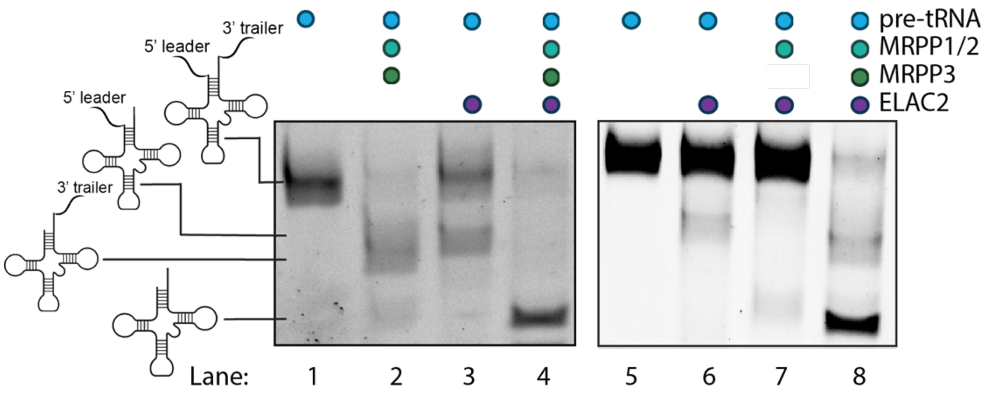


**Figure S3 | Wild-type pre-tRNA$^{Tyr}$ cleavage in the presence different protein combinations |** *In vitro* cleavage assay of pre-tRNA$^{Tyr}$ with its 5′ leader and 3′ trailer. RNA was resolved on a 7% urea-polyacrylamide gel, stained with 1x SYBR Gold, gel representative of 3-5 replicates. Each circle at the top indicates the presence of the specified reaction component, i.e., pre-tRNA substrate or components of the RNase complexes.

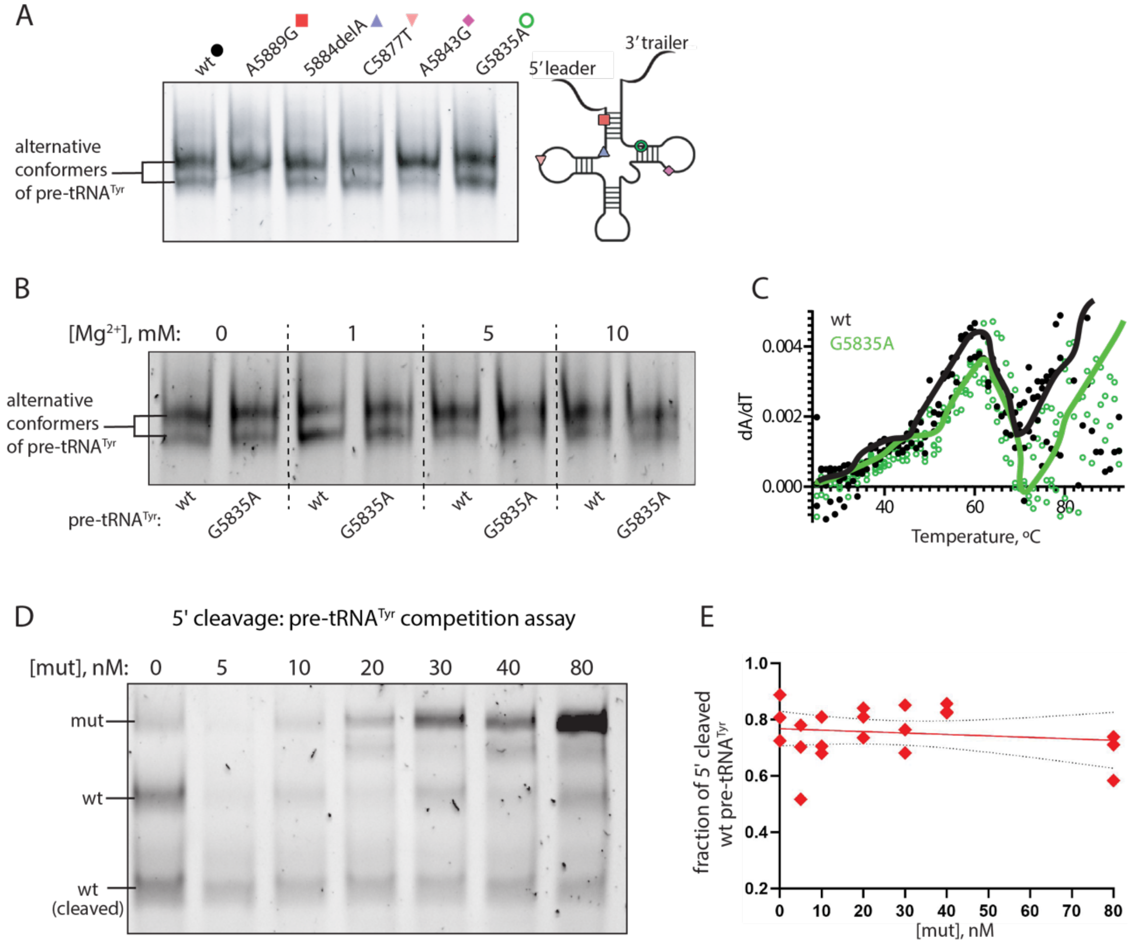


**Figure S4 | Variant effects on tRNA conformation and competitive inhibition of 5′ leader removal | (A)** 8.5% native-PAGE of pre-tRNA$^{Tyr}$ substrates containing the 5′ leader and 3′ trailer. Two bands suggest two alternate conformers of the RNA. Stained with 1x SYBR Green II. **(B)** 8.5% native-PAGE of the pre-tRNA$^{Tyr}$ as a function of $Mg^{2+}$ concentration during refolding. **(C)** UV melting profile for the wt and G5825A pre-tRNA$^{Tyr}$ containing the 5′ leader and 3′ trailer in 1 mM $Mg^{2+}$ refolding buffer. Three replicates for each sample are plotted with a representative melting profile curve of the average plot. **(D)** *In vitro* 5′ cleavage assay of wt pre-tRNA$^{Tyr}$ (“wt” on the gel) premixed with the A5843G mutant of pre-tRNA$^{Tyr}$-tRNA$^{Cys}$ ( “mut” on the gel) prior to protein introduction. The mut was titrated (0-80 nM final concentration) vs the wt was kept at a constant 40 nM. Samples were run on a 7% urea-polyacrylamide gel and stained with 1x SYBR Gold. Gel representative of three biological replicates. **(E)** Graph showing the fraction of 5′ processed wt pre-tRNA$^{Tyr}$ in the presence of the different A5843G mutant concentrations. The trendline (red) with a slope of -0.0005 and 95% confidence integral (black dotted lines) of -0.002 to 0.001. Measured from Fig. S4D gel, quantified using ImageLab (BioRad) and plotted using GraphPad prism.

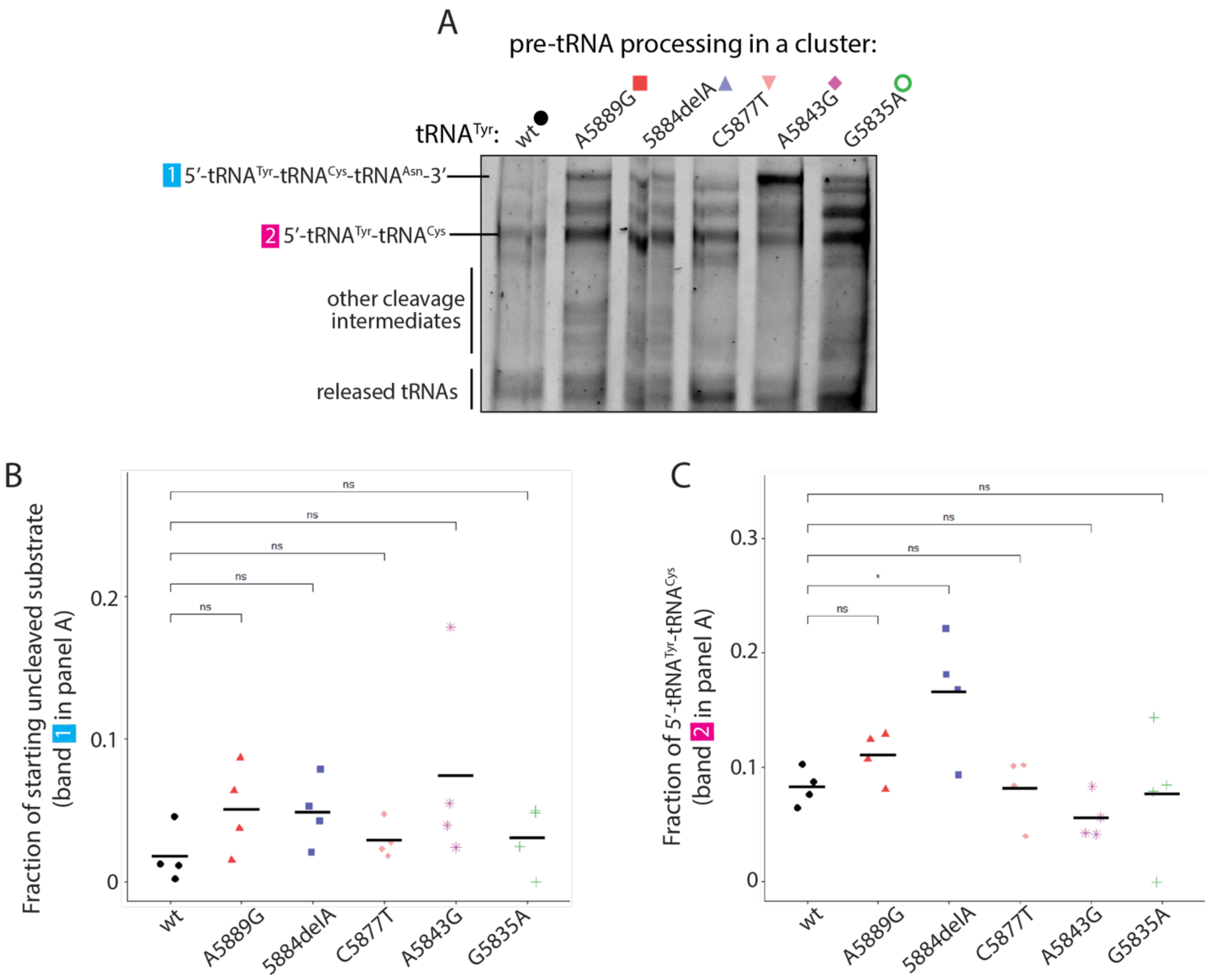


**Figure S5 | Release of tRNA$^{Asn}$ from the tRNA$^{Tyr}$-tRNA$^{Cys}$-tRNA$^{Asn}$ cluster is unaffected by the tRNA$^{Tyr}$ variants | (A)** *In vitro* cleavage gel showing concurrent processing of the pre-tRNA$^{Tyr}$-tRNA$^{Cys}$-tRNA$^{Asn}$ substrate with RNase P and RNase Z. The RNA species were resolved on a 7% urea-polyacrylamide gel and stained with 1x SYBR Gold. The gel is representative of four biological replicates. The tRNA$^{Tyr}$-tRNA$^{Cys}$ cleavage product (band 2) was identified with a pre-tRNA$^{Tyr}$-tRNA$^{Cys}$ substrate loaded alongside the cleavage reactions (not shown) to demonstrate that tRNA$^{Asn}$ is being processed successfully from the polycistronic transcript across the sequence variant substrates. **(B)** and **(C)** quantitation of bands 1 and 2 using ImageLab (BioRad). Data plotted using R. A one-way ANOVA was used for statistical analysis.

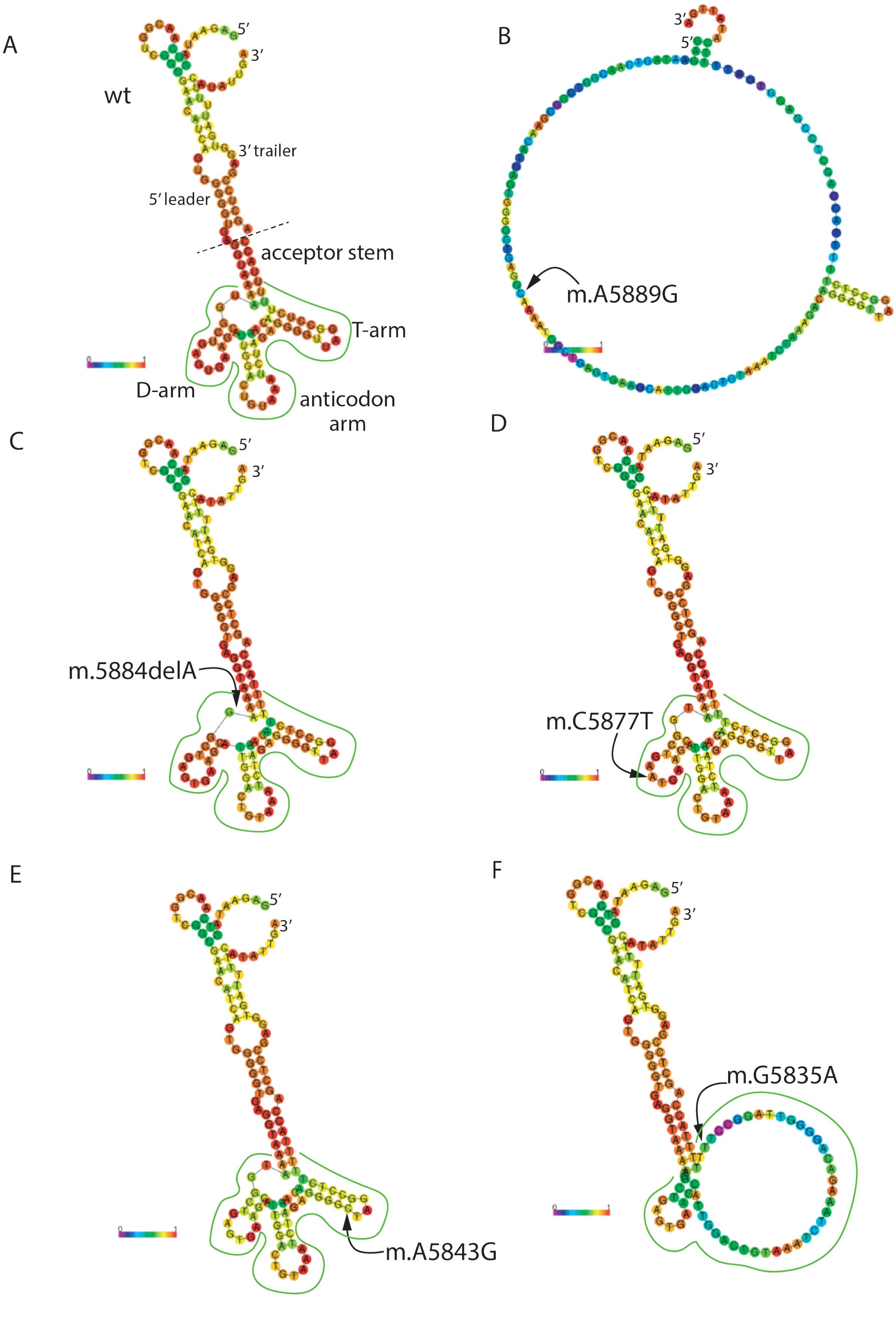
A
wt
5′
3′
3′ trailer
5′ leader
acceptor stem
T-arm
D-arm
anticodon arm
B
3′
5′
m.A5889G
C
5′
3′
m.5884delA
D
5′
3′
m.C5877T
E
5′
3′
m.A5843G
F
5′
3′
m.G5835A

**Figure S6 | RNAfold centroid structure models of the wild-type and variant tRNA$^{Tyr}$ |** Predicted secondary structures of pre-tRNA$^{Tyr}$ containing its native 5′ 38-nt leader and 24-nt 3′ trailer sequences. The tRNA$^{Tyr}$ cloverleaf secondary structure region is outlined in green for all models except panel B where base pairing between the leader and trailer appears to be majorly disrupted. The centroid structure represents the average of all potential structures with the coloring for base-pair probabilities (0 = pink, red = 1).

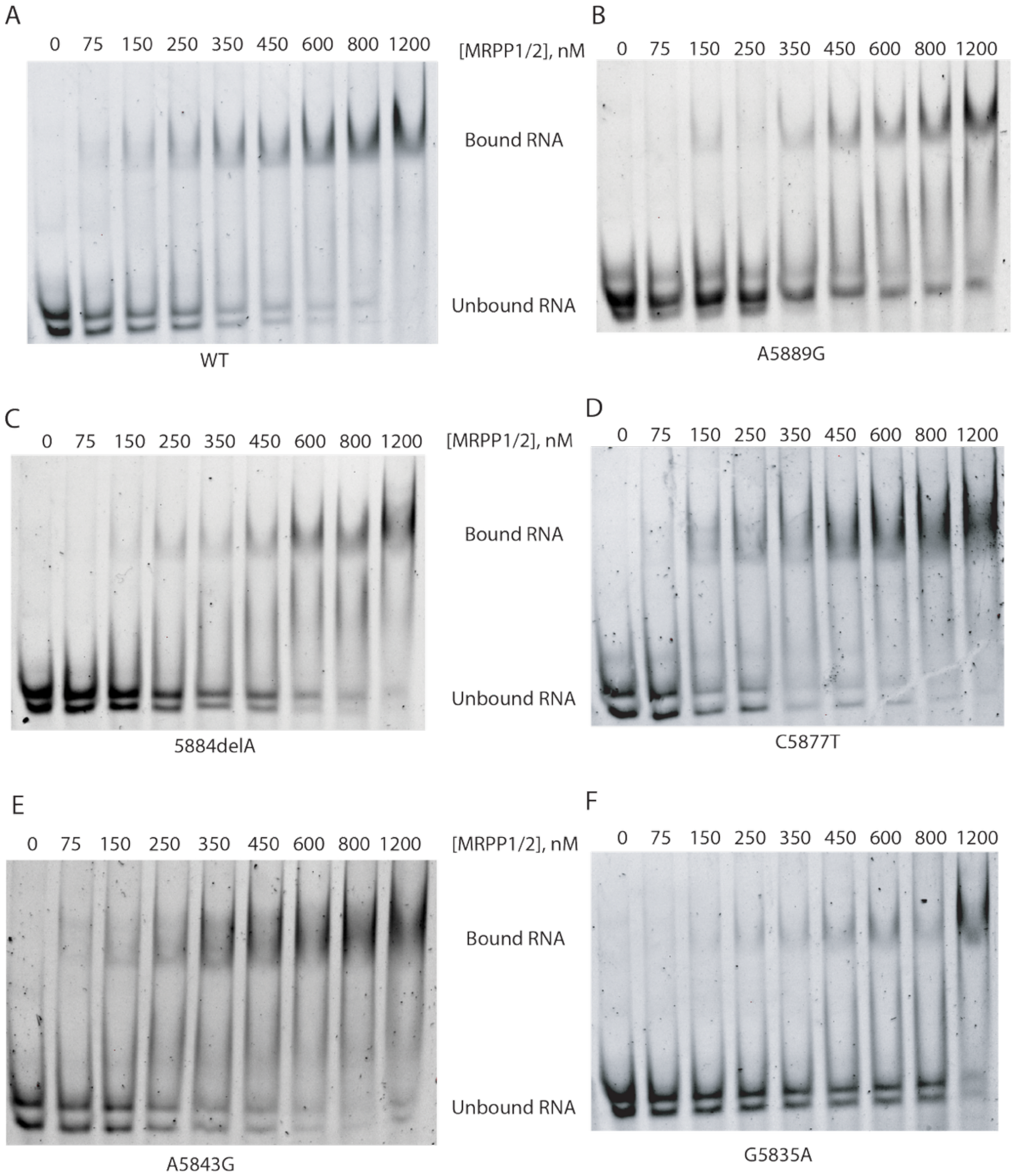


**Figure S7 | Representative EMSA gels for wt and variant pre-tRNA$^{Tyr}$ |** Electromobility Shift Assay (EMSA) gels to analyze binding of pre-tRNA$^{Tyr}$ containing its native 5′ 38-nt leader and 24-nt 3′ trailer with an increasing concentration of MRPP1/2. Samples resolved on 8.5% native polyacrylamide gels and stained with SYBR Green II. Each gel representative of four biological replicates.

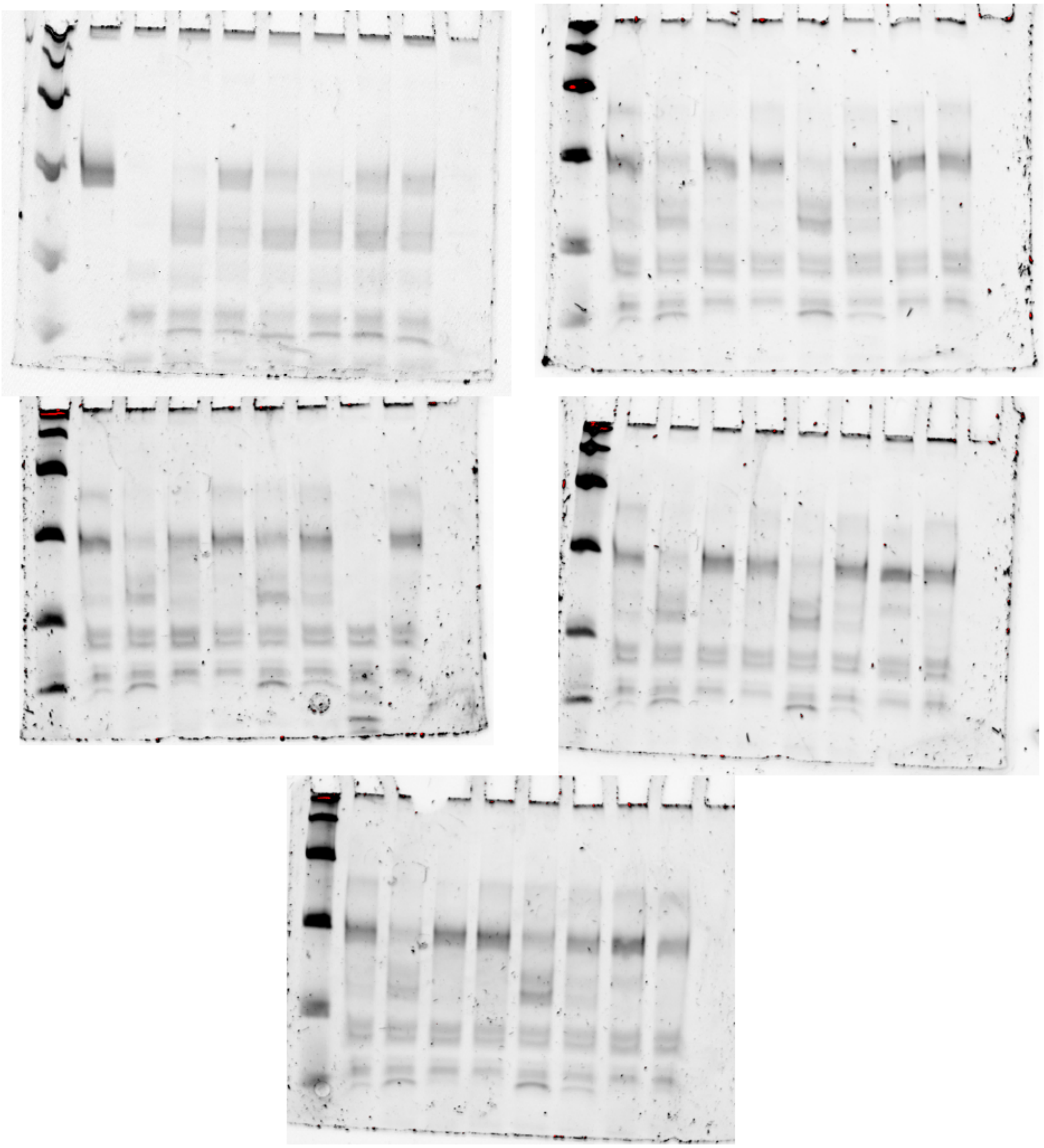

**Figure S8 | Fig 2A full gel images |**

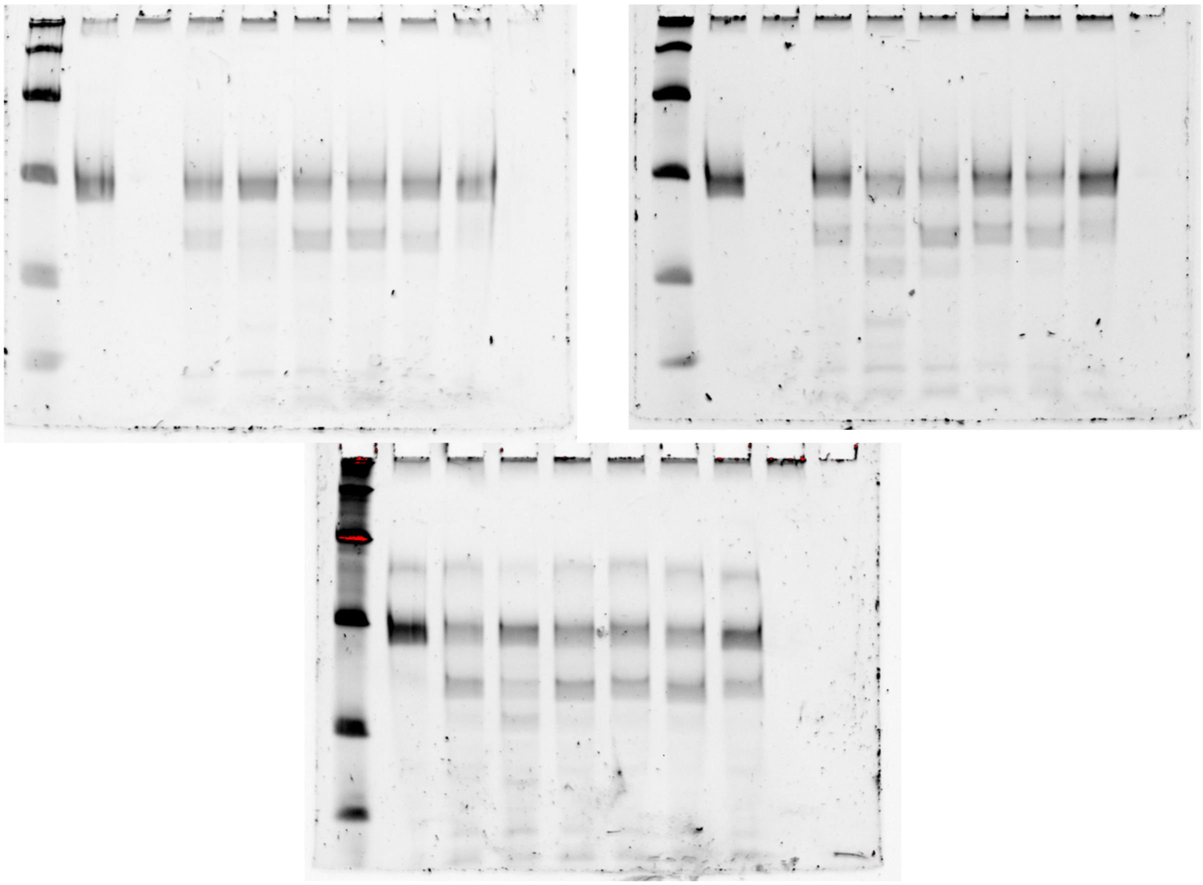

**Figure S9 | Fig 2B full gel images |**

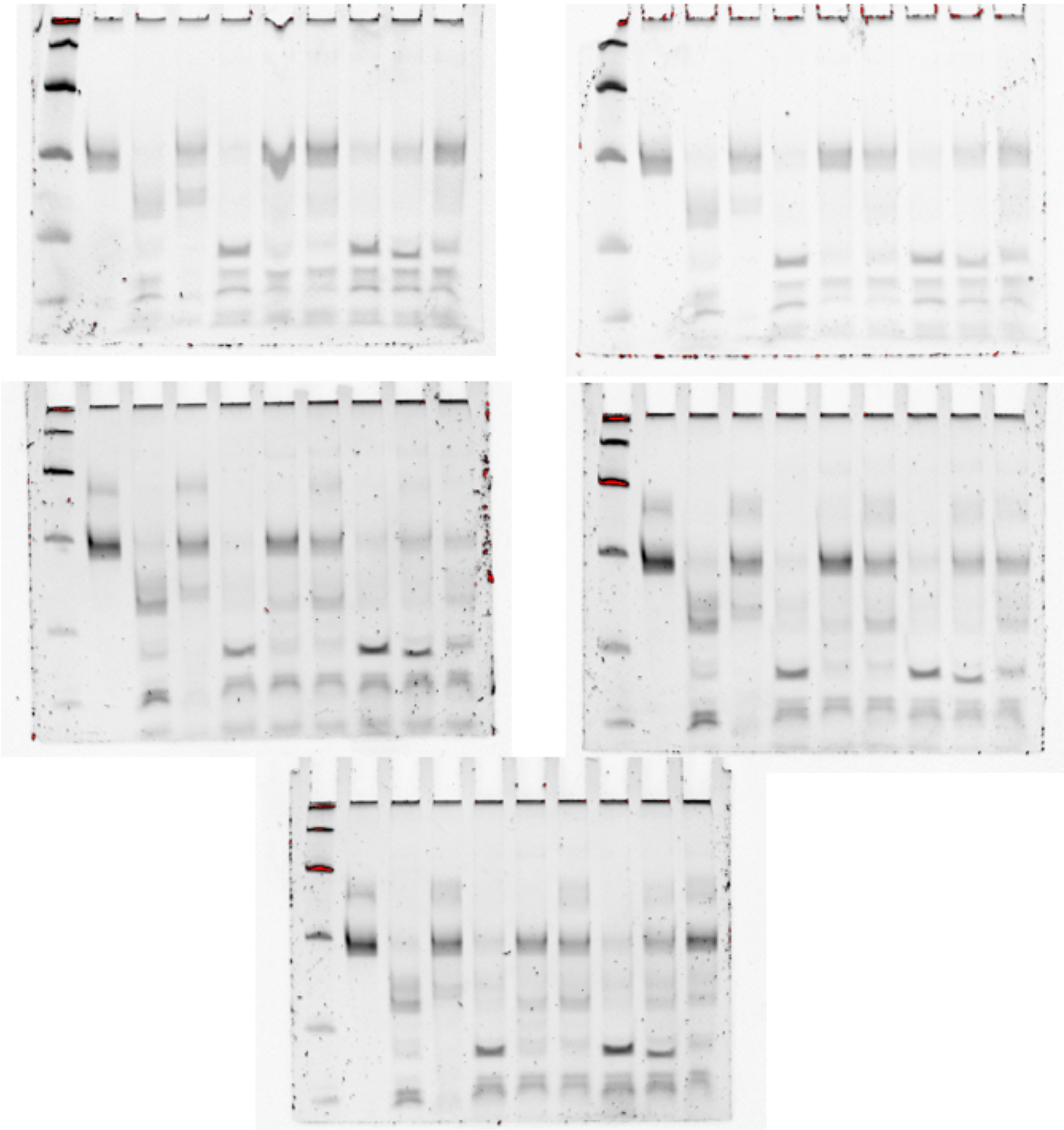

**Figure S10 | Fig 3 full gel images |**

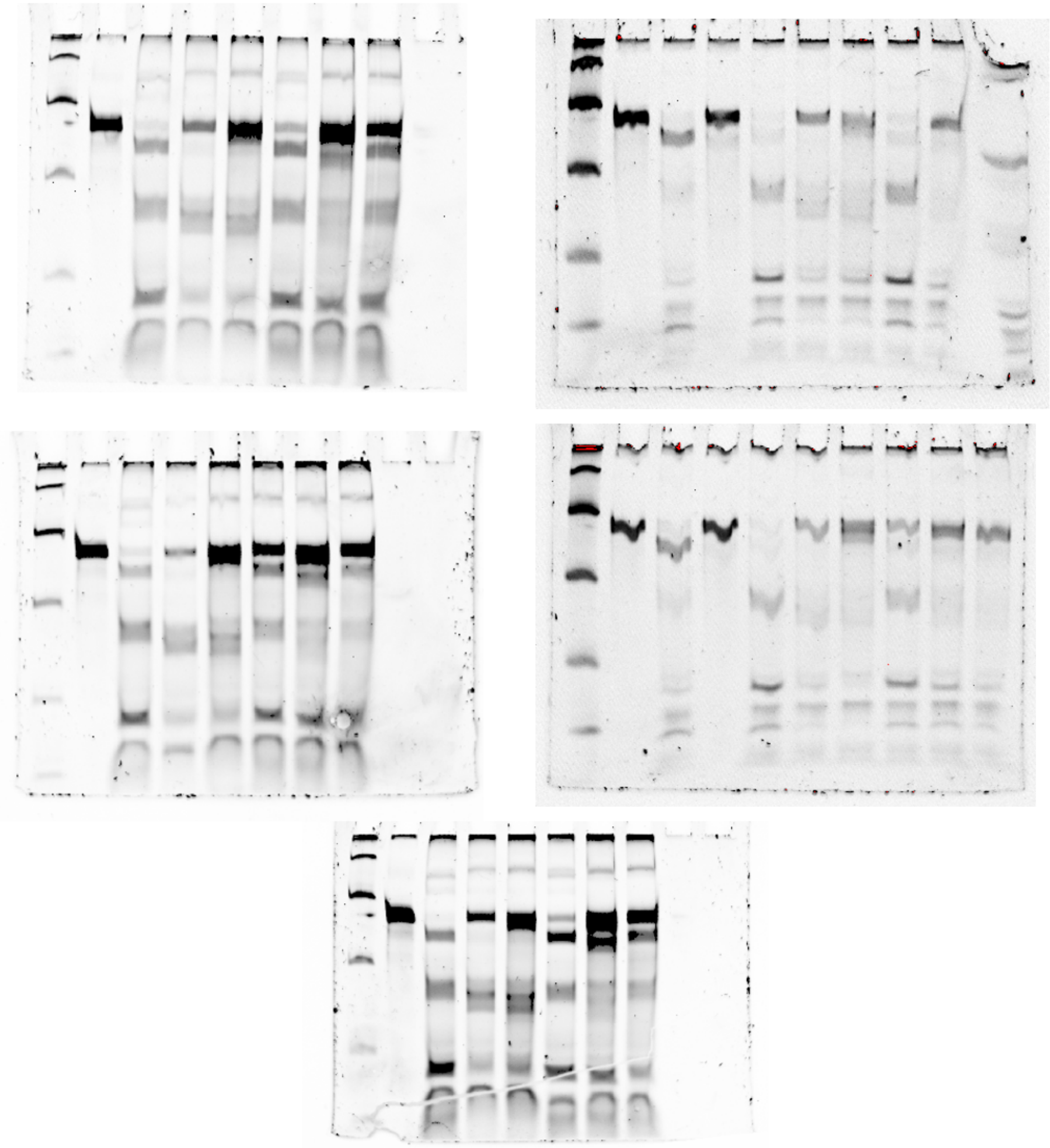

**Figure S11 | Fig 5 full gel images |**
